# Diversification of fungal chitinases and their functional differentiation in *Histoplasma capsulatum*

**DOI:** 10.1101/2020.06.09.137125

**Authors:** Kristie D. Goughenour, Janice Whalin, Jason C. Slot, Chad A. Rappleye

## Abstract

Chitinases enzymatically hydrolyze chitin, a highly abundant biomolecule with many potential industrial and medical uses in addition to their natural biological roles. Fungi are a rich source of chitinases, however the phylogenetic and functional diversity of fungal chitinases are not well understood. We surveyed fungal chitinases from 373 publicly available genomes, characterized domain architecture, and conducted phylogenetic analyses of the glycoside hydrolase family 18 (GH18) domain. This large-scale analysis does not support the previous division of fungal chitinases into three major clades (A, B, C). The chitinases previously assigned to the “C” clade are not resolved as distinct from the “A” clade in this larger phylogenetic analysis. Fungal chitinase diversity was partly shaped by horizontal gene transfer, and at least one clade of bacterial origin occurs among chitinases previously assigned to the “B” clade. Furthermore, chitin binding domains (CBD) including the LysM domain do not define specific clades but instead are found more broadly across clades of chitinase enzymes. To gain insight into biological function diversity, we characterized all eight chitinases (Cts) from the thermally dimorphic fungus, *Histoplasma capsulatum:* six A clade (3 A-V, 1 A-IV, and two A-II), one B clade (B-I), and one formerly classified C clade (C-I) chitinases. Expression analyses showed variable induction of chitinase genes in the presence of chitin but preferential expression of *CTS3* in the mycelial stage. Activity assays demonstrated that Cts1 (B-I), Cts2 (A-V), Cts3 (A-V), Cts4 (A-V) have endochitinase activities with varying degrees of chitobiosidase function. Cts6 (C-I) has activity consistent with N-acetyl-glucosaminidase exochitinase function and Cts8 (A-II) has chitobiase activity. This suggests chitinase activity is variable even within sub-clades and that predictions of functionality require more sophisticated models.

## INTRODUCTION

Chitin is a (1,4)-β-linked N-acetyl-D-glucosamine (GlcNAc) polymer. As the second most abundant biopolymer after cellulose (Tharanathan and Kittur 2003), chitin and its deacetylated derivative chitosan are abundant sources of organic carbon and nitrogen that have many potential industrial uses, from biomedical to agricultural to water engineering (Ravi Kumar 2000; Zargar et al. 2015). Consequently, there is a great interest in identifying enzymes that efficiently hydrolyze chitin into more soluble mono- and oligomers of GlcNAc. Chitin degrading enzymes also have potential applications in the breakdown of the chitinous structures of fungal and arthropod agricultural pests (Hamid et al. 2013).

Chitinases (E.C 3.2.1.14) are glycosyl hydrolases that are found in a wide range of plants, bacteria, and fungi. For plants and bacteria, which lack chitin, chitinases play roles primarily in defense against fungi and/or arthropods. Chitin is an important structural component of the fungal cell wall, ranging from 0.5%-5% in yeasts to ≥20% in some filamentous fungi (Hartl et al. 2012). Yeast-form fungi possess relatively few chitinases (e.g., two in *Saccharomyces cerevisiae*) compared to the 10-20 chitinases encoded in some filamentous fungal genomes (Seidl 2008). The expansion of chitinases in filamentous fungi has prompted investigation of chitinase roles in formation of hyphal structures (Seidl 2008). While some studies have shown select chitinases are specifically induced during hyphal formation (Gruber, Kubicek, and Seidl-Seiboth 2011; Takaya et al. 1998), many other chitinases are not (Duo-Chuan 2006; Gruber, Vaaje-Kolstad, et al. 2011; Seidl 2008) suggesting alternative roles for diverse chitinases. For example, there is evidence that specific chitinases facilitate mycoparasitism of other fungi by *Trichoderma* (Boer et al. 2007; Cruz et al. 1992; Seidl et al. 2005).

Fungal chitinases are characterized by the presence of the gycoside hydrolase 18 family (GH18) (Seidl 2008) domain. Additional motifs often found in chitinases include: an N-terminal signal peptide region that serves for secretion of the enzyme, a serine/theronine-rich region, one or more chitin-binding domains (CBDs), and LysM domains. The LysM domain enables binding to polysaccharides such as peptidoglycan and chitin (i.e. a CBD). However, none of these motifs beyond the GH18 domain are conserved across all fungal chitinases or required for a protein to be considered a chitinase (Duo-Chuan 2006).

A three-clade classification system for fungal chitinases (clades A, B, and C) emerged from several investigations into the diversity of GH18-domain containing enzymes among fungi (Karlsson and Stenlid 2008, 2009; Seidl 2008; Seidl et al. 2005). While some studies have further divided the B clade into D and E clades, this appears to be a result of their specific datasets rather than a widely applicable feature (Junges et al. 2014). Clade A chitinases are between 40-50 kDa and were previously reported to have no CBD. These are the most well-studied of the three clades (Seidl 2008). Clade B chitinases are variable in size and also in the presence and number of CBDs. Clade C chitinases are distinguished by a significantly larger size (140-170 kDa) due to extension at the C-terminus. Clade C chitinases were also proposed to be defined by the presence of multiple CBDs, and especially LysM motifs (Seidl et al. 2005; Karlsson and Stenlid 2008; Karlsson and Stenlid 2009). Clade C chitinases are the least well characterized of all the clades; few members have been included in previous phylogenetic studies, and none have been functionally characterized (Duo-Chuan 2006; Seidl 2008). However, this three clade classification of fungal chitinases is based on a limited number of fungal genomes, and it remains to be determined if it is robust to more in-depth taxon sampling (Seidl et al. 2005; Karlsson and Stenlid 2008; Karlsson and Stenlid 2009).

While many fungal chitinases have been transcriptionally profiled, the characterization of fungal chitinase enzymatic specificity is particularly limited, leaving broad assumptions about clade-specific functions untested. For example, Clades A and C are class V (fungal/bacterial) chitinases, which are generally assumed to be exochitinases based on modeling of the conserved binding groove as deep and tunnel shaped (Duo-Chuan 2006; Seidl 2008; Hartl et al. 2012). Clade B corresponds to class III (fungal/plant) chitinases and are predicted to be endochitinases due to the modeling of their binding grooves as shallow and open (Duo-Chuan 2006; Seidl 2008; Hartl et al. 2012). While some of these assumptions are supported by the activities of specific chitinases (i.e., the CHIT33 and CHIT42 chitinases of *Trichoderma harzianum* (Boer et al. 2007; Lienemann et al. 2009)), these chitinases also have multiple activities and further examples need to be studied to establish clade-defning characteristics (Hartl et al. 2012). The inconsistent use of diverse chitin substrates (e.g., crustacean chitin or fungal chitin) further confounds the reliability of assumptions about functional diversity within and among clades. While there are reports of complete chitinase or clade/sub-clade specific chitinase transcriptional or deletion studies (Alcazar-Fuoli et al. 2011; Dünkler et al. 2005; Gruber, Vaaje-Kolstad, et al. 2011; Gruber, Kubicek, et al. 2011), enzymatic studies have not been as comprehensive.

In this study, we identified the GH18 domain proteins encoded in 373 published fungal genomes to conduct a more complete analysis of their distribution and evolution. In addition, we provide initial expression analysis and enzymatic characterization of the chitinase enzymes produced by the thermally dimorphic fungus, *Histoplasma capsulatum*. This organism has distinct and tightly controlled morphologies (mycelia or yeasts), each of which has a specific lifestyle (environmental saprobe or pathogen of mammals, respectively) potentially allowing for identification of specific roles for different chitinases. These chitinases provide new enzymatic examples from each of the major clades, including the enzymatic activity of the first characterized clade C chitinase.

## RESULTS

### Diversification of fungal GH18 domains

We identified 3,888 GH18 domain-containing proteins in 373 publicly available fungal genomes (Supplemental Data 1) using an HMM search (Eddy 2009). 494 of these proteins contain the LysM chitin binding domain (Supplemental Data 2) considered to be characteristic of the C clade chitinases in fungi (Seidl et al. 2005; Seidl 2008). 1,250 CBDs (i.e., ChtBD1, ChtBD3, COG3979, CBM_1 and CBM_19) were also identified in the GH18 domain containing proteins (Marchler-Bauer et al. 2017).

Alignment of the GH18 domains of these proteins was 1,840-characters long; however, the C-terminus of the alignment is not well-conserved and occasionally absent particularly among members of the B clade (including functionally validated chitinases (Dünkler et al. 2005; Hurtado-Guerrero and van Aalten 2007)). Maximum likelihood phylogenetic analysis using IQ-Tree was used to establish evolutionary relationships among the fungal chitinases and to infer support for clades and subclades defined by the common ancestor of previously classified chitinases, with some slight expansion for closely related but previously unstudied taxa (Figure 1, Table 1, Supplemental Data 3). This identified the previously reported A+C clade, although this was not well-supported (70% of rapid bootstraps (RB)). The B clade was not supported. Constituent sub-clades, determined by identifying the most recent common ancestor of previously categorized chitinase sub-clades (Seidl et al. 2005; Karlsson and Stenlid 2008; Karlsson and Stenlid 2009), were generally recovered with variable support. Further expansion of sub-clades recovers more strongly supported nodes without encroaching on other sub-clades (Figure 1).

**Figure 1.**
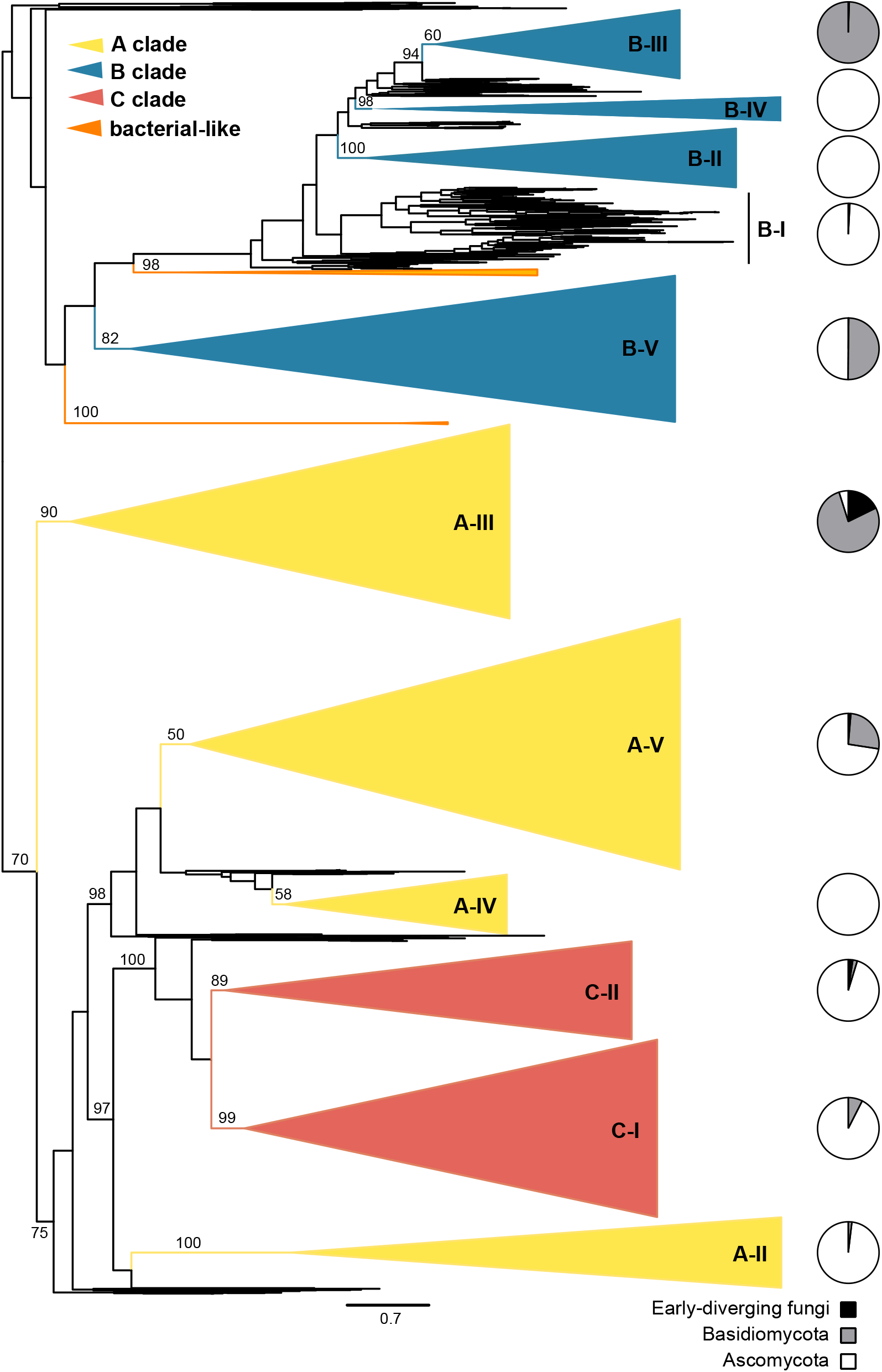
Phylogenetic analysis of fungal GH18 domains. Phylogenetic tree depicts relationships among 3,888 fungal chitinase proteins based on alignment of their glycosyl hydrolase (GH18) domains. Trees were built using maximum-likelihood methods (IQ-Tree) and branch support assessed by 1000 Ultrafast bootstrap replicates (values indicated by numbers at the root of common ancestor branches). Major groups corresponding to the previously named A, B, and C clades are indicated by color (yellow, blue, red, respectively). Clades comprised of bacterial-like GH18 proteins are indicated (orange). Collapsed sub-clades, indicated as triangles with names to the right, are defined by the most recent common ancestor of previously categorized chitinases, in some cases extended to supported clades consistent with the fungal phylogeny. The tree is rooted along the midpoint, which generates the B vs AC clade split that has been widely accepted in the literature (Karlsson and Stenlid 2008, 2009; Seidl et al. 2005). Pie charts at the far right show the fraction of the constituent chitinase proteins in each clade occurring in early-diverging fungi (black), Basidiomycota (gray), and Ascomycota (white).

**Table 1.**
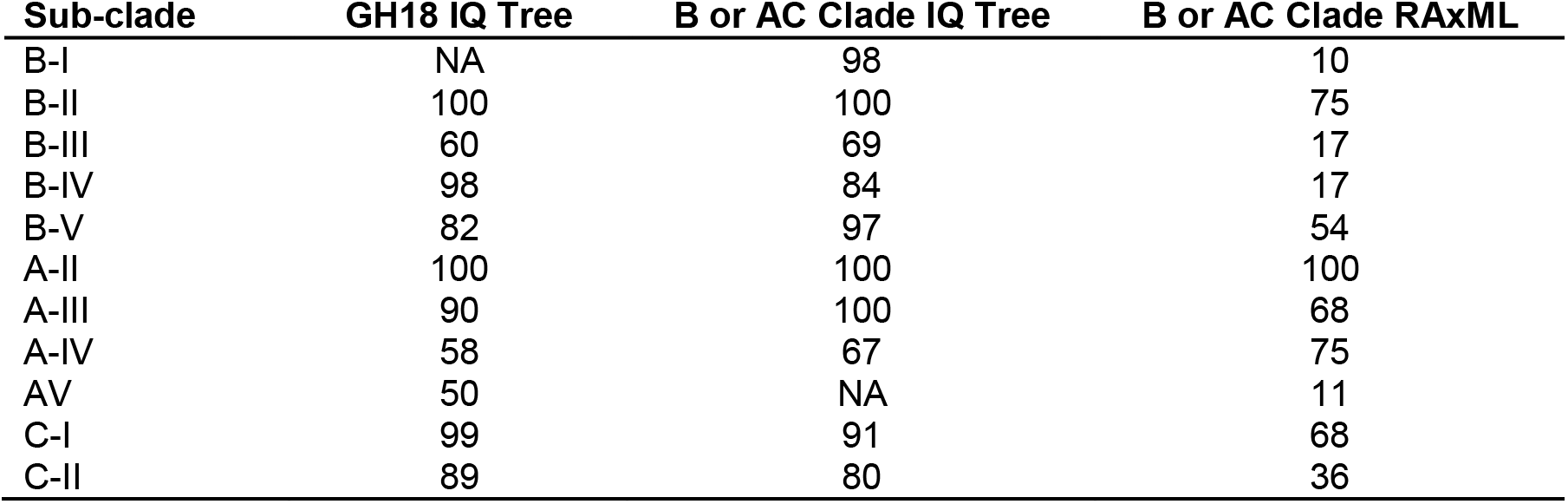
Bootstrap support values for sub-clades by various phylogenetic analyses.

In order to retain more alignable characters and thereby reduce error introduced by homoplasy for finer resolution within clades and sub-clades, we separately analyzed sequences on either half of a bifurcation between the A+C clades and an assemblage containing all the B sub-clades in the complete tree (Figure 2). In the complete GH18 tree (Figure 1), we did not recover a B-I sub-clade and only the B-II and B-IV sub-clades received strong rapid bootstrap (RB) support (Table 1). However, in the IQ-tree focused on B sub-clades, B-I (98% RB), B-II (100% RB), and B-V (97% RB) clades were supported, while clades B-III and B-IV were not supported with the current strict definition of sub-clades, and a B clade remained unsupported (Figure 2A and Table 1). However, slight expansions of the sub-clades identified supported nodes consistent with the previous analyses. A RAxML analysis of this alignment also recovered these B sub-clades but without RB support, suggesting the finer scale topology is not always robust to differences in methodology (Table 1 and Supplemental Data 3).

**Figure 2.**
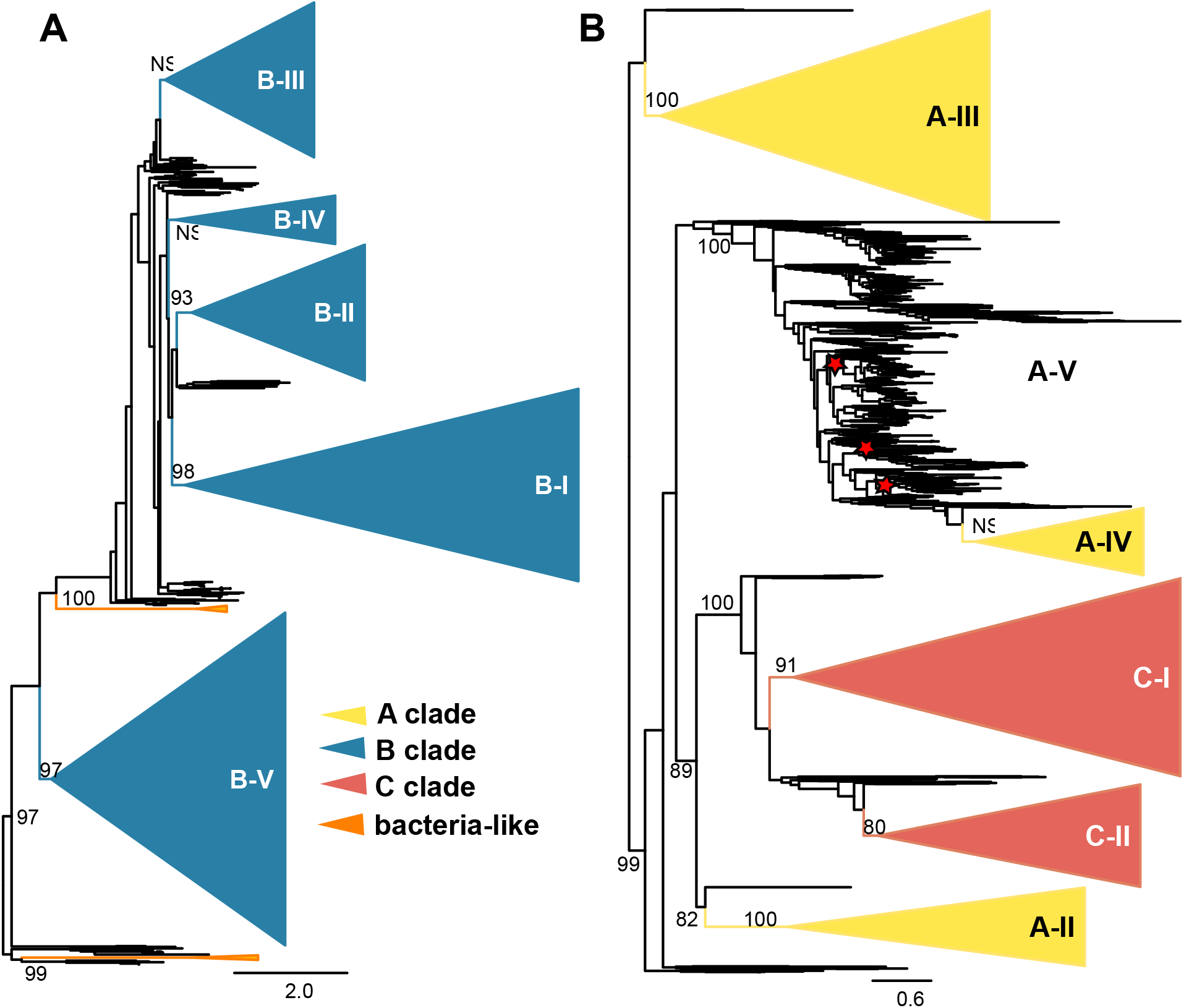
Phylogenetic analysis of fungal B and AC clades. Relationships among constituent proteins in the B and AC clades when analyzed as separate groups. Trees were built using maximum-likelihood methods (IQ-Tree) and branch support assessed by 1000 Ultrafast bootstrap replicates (values indicated as percentages at the base of the clade). Trees are rooted as in the complete GH18 IQ-Tree in Figure 1. Collapsed sub-clades, indicated as triangles with names to the right, are defined by the most recent common ancestor of previously categorized chitinases, in some cases extended to supported clades consistent with the fungal phylogeny. (**A**) IQ-Tree based phylogeny of B clade chitinase proteins with sub-clades indicated and collapsed (blue). (**B**) IQ-Tree based phylogeny of chtinases belonging to the A (yellow) and C clades (red) showing the evolution of the C clade as a branch of the A clade. As sub-clade A-V is polyphyletic in the AC tree, the ancestral nodes giving rise to previously identified chitinases are indicated with red stars.

For the branch in the entire GH18 tree containing the A and C clades (hereafter referred to as the “AC” clade), we recovered distinct, yet unsupported A-IV (58% RB) and A-V (50% RB) sub-clades after reclassifying one *Trichoderma reesei* gene (Chi18-5) from an A-V to an A-IV (Figure 1 and Table 1). In the AC-only IQ-Tree (Figure 2B), the A-V sub-clade was not monophyletic. The A-II (100% RB, IQ-Tree) and A-III (100% RB, IQ-Tree) sub-clades were supported in all analyses (Figure 1, Figure 2B, and Table 1). The A-IV and A-V sub-clades were separated (98%RB, IQ-Tree) from the A-II and C clades (Figure 1). While the C group is monophyletic within the AC clade (Figure 1), separation into sub-clade C-I was supported (99% RB) while sub-clade C-II was not (89% RB) in the complete tree (Figure 1, Table 1, Supplemental Data 3). In the separate AC tree C-II is not supported but would be supported by including sister sequences (Figure 2B). Interestingly, in our analysis clade C (both C-I and C-II) groups closely, with strong support, with A-II (97% RB in the all GH-18 IQ-Tree) to the exclusion of other A sub-clades (Figure 1 and Supplemental Data 3).

### Taxonomic distribution of GH18 chitinases

Each sub-clade hosts a distinct assortment of fungal taxa. The Ascomycota, particularly Leotiomycetes, Sordariomycetes, Eurotiomycetes, and Dothideomycetes, have the largest number of chitinases and highest diversity of chitinase sub-clades (Figure 3, Supplemental Data 4). B sub-clades-I, -II, and -IV are composed almost entirely of Ascomycota. The individual orders Malasseziales, Glomerales, Monoblepharidales, Neocallimastigales, and Rozella have the fewest chitinases per genome (e.g. 1 or 0.5 (Rozella)) and Auriculariales, Basidiobolales, Geastrales, and Xylariales have the most (greater than 17). The B-IV sub-clade consists entirely of Saccharomycetales, while the B-I and B-II sub-clades are composed of non-Saccharomycetales, Ascomycota. B-I consists of Leotiomycetes, Agaricomycetes, Dothideomycetes, Eurotiomycetes, Leotiomycetes, Sordariomycetes, and Xylonomycetes, with multiple orders of Dothideomycetes, Eurotiomycetes and Sordariomycetes represented. B-II contains multiple orders of Eurotiomycetes, Dothideomycetes, Leotiomycetes, and Sordariomycetes. Sub-clade B-V contains a large number of Basidiomycota, however several species of Ascomycota and a single early diverging fungus (*Basidiobolus meristosporus*) are also represented. B-III is composed almost entirely of Basidiomycota with the exception of sequences from the early diverging Mucorales.

**Figure 3.**
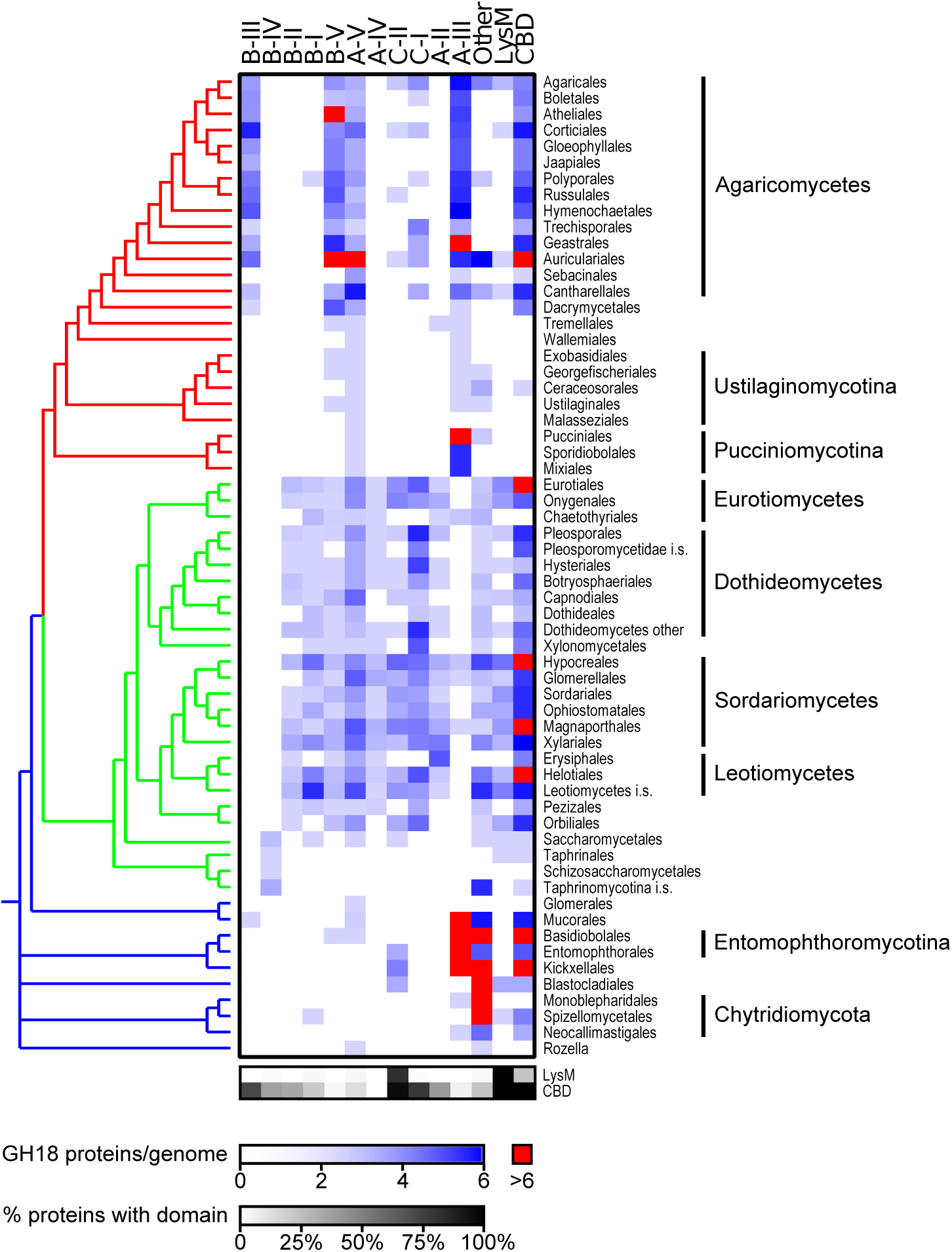
Distribution of chitinases among fungal taxa. Heat map showing the distribution of chitinase sub-clades across fungal orders. Shading indicates the average number of GH18 domain-containing proteins belonging to each sub-clade (columns) per genome with orders with highly expanded numbers (at least 6 per genome) of sub-clade chitinases indicated (red). Taxonomic relationships among fungal orders is presented for reference (left side) and colored for early-diverging fungi (blue), Ascomycota (green), and Basidiomycota (red). The right two columns of the heat map indicate the number of chitinases per genome with LysM or CBD domains, respectively. Lower grayscale heat map indicates the percentage of the subclade proteins with LysM and CBD domains.

Within the AC clade, sub-clades A-II and A-IV are composed mostly of Ascomycota with isolated Basidiomycota (Tremellales in A-II and Pucciniales in A-IV) (Figure 3). The A-II sub-clade consists of Eurotiomycetes, Sordariomycetes, Leotiomycetes, and Dothideomycetes, although within the Dothideomycetes the order Capnodiales and an unnamed order in Pleosporomycetidae were not represented. Only the Dothideales, Orbiliales, and Lecanorales were not represented in A-IV. A-III is composed of both Basidiomycota and early diverging fungal chitinases with a few Ascomycota (Sordariomycetes). Malasseziales was the only order of Basidiomycota with no A-III chitinases while in the early diverging fungi Glomerales, Blastocladiales, and Rozella also lacked these chitinases. The A-V sub-clade contains Ascomycota, Basidiomycota and early diverging fungi. All Ascomycota and Basidiomycota were represented in this clade except one member of an unnamed class in Taphrinomycotina, and among early diverging fungi, they are only found in Rozella, Mucorales, Glomerales, and Basidiobolales. The C-I sub-clade mostly contains Ascomycota with a small assortment of sequences from 6 orders of Agaricomycetes while the C-II sub-clade contains both Ascomycota and a few early diverging fungi (Kickxellales, Entomophthorales, and Blastocladiales). Relatively few chitinases are found in Botryosphaeriales.

### Patterns of GH18 chitinase diversification

Much of the GH18 chitinase phylogeny is consistent with vertical inheritance. More derived branches of the chitinase phylogeny frequently track the species phylogeny, particularly in the A sub-clades (A-V, A-IV, A-III, and A-II) and the B-V sub-clade. In keeping with the fungal phylogeny, the B-V sub-clade contains distinct Agaricomycetes, Eurotiomycetes, and Dothideomycetes groups, and a single species of Leotiomycetes within a Sordariomycetes group. Similarly, the A-IV sub-clade contains distinct clades of Sordariomycetes, Eurotiomycetes, Leotiomycetes, and Dothideomycetes. The A-II sub-clade has large separate Sordariomycetes and Dothideomycetes groups. The A-V sub-clade contains a distinct Saccharomycetes clade, but there are also multiple Agaricomycetes, Sordariomycetes, and Eurotiomycetes clades. There are large-scale divergences from the fungal tree, including a clade of Sordariomycetes within a clade otherwise composed of Agaricomycetes in the A-III sub-clade. Despite such grouping of related species, the branching order within GH18 phylogenies is overall complex, suggesting domain and gene duplications are common. Some sub-clades in particular (e.g. the C sub-clades) show few taxon-specific clades (Figure 3).

Part of the contrast with the species tree comes from specific supported instances of horizontal gene transfer (HGT) (Supplemental Figure S1). For example, two clades of mainly Hypocreales (Pezizomycotina) fungi were initially poorly-placed among the B sub-clades in the GH18 phylogenies. A re-analysis including fungal and non-fungal sequences at NCBI more confidently placed these sequences among bacterial chitinases (Figure 1), suggesting the clades originated through HGT, further reducing support for a B clade. One of these chitinases appears to have been acquired specifically by insect-pathogenic Hypocreales fungi (e.g. XP_006673222.1 *Cordyceps militaris*) from bacteria of an unknown lineage (Supplemental Figure S1A). This *C. militaris* chitinase is an endo-N-acetylhexosaminidase active on fucose-containing N-glycans (Huang et al. 2018). The other chitinase, found in a much larger group of fungi in Hypocreales and additional fungi in Pezizomycotina is supported to have been transferred from Actinobacteria to Hypocreales (e.g. XP_006670951.1 *Cordyceps militaris*, Supplemental Figure S1B) and later to other fungi. This single-domain chitinase is most similar to chitinase D (cd02871), which has hydrolytic and transglycosylation activity in bacteria (Vaikuntapu et al. 2016). Two other HGTs of chitinase D were supported (Supplementary Figure S1C), one from Streptomyces (Actinobacteria) to Arthrodermataceae (Eurotiomycetes), and the other from Micromonadaceae (Actinobacteria) to Auriculariales (Agaricomycetes). (Vaikuntapu et al. 2016). Still another group of insect-associated Hypocreales chitinases (e.g. KOM21536.1 *Ophiocordyceps unilateralis*, Supplemental Figure S1D) is dispersed in a clade of fungal chitinases otherwise composed of multiple paralogs from the arthropod-associated Kickxellomycotina. These chitinases are part of the expanded A-III sub-clade, and contain a GH18 domain and a signal secretion peptide. Outside Hypocreales, the amphibian gut symbiont *Basidiobolus meristosporus* appears to have acquired a B-V chitinase with only a GH18 domain (Basme2_176417, Supplemental Figure S1E) from Agaricomycetes, possibly in Auriculariales, but a specific donor is not supported. Finally, another inter-phylum transfer appears to have occurred from Pezizomycotina to Panaeolus (Pancy2_12872, Agaricales, Supplemental Figure S1F). The nearest sequence, in *Uncinocarpus reesei* (Uncre1_6593), associates this last event with the dung decay niche. This is a C-II chitinase with 2 LysM and 1 CBD domain. Analyses comparing topological constraints were able to reject monophyly of the hypothetical donor groups for most, but not all HGTs, due to the complexity of the overall phylogeny and an absence of a reliable outgroup in some cases (Supplementary Data 5).

### Evolution of domain architecture

Chitinases with multiple GH18 domains were rare, observed in only 11 cases. Of these, all contained two GH18 domains except *Psilocybe cyanescens* [Psicy2_12260], which contained three. All multi-GH18 domain proteins were in Basidiomycota; most were in Agaricales (9 of 11 cases) and there was one each in Polyporales and Pucciniales. Multi-GH18 domain proteins are not constrained to a specific sub-clade, and are found in the A-III (6 cases), B-III (3 cases) and B-V (2 cases) sub-clades. Interestingly only 5 GH18 domains in multi-GH18-containing proteins, all in B sub-clades, were predicted to be active. In addition, both GH18 domains in multi-domain A-III chitinases lacked conserved active site residues.

LysM domains occur in 4 chitinase sub-clades. Most are restricted to the C-II sub-clade (Figure 3 and Supplemental Data 2), which also includes the previously described LysM containing class III Hce2 (Homologs of *C. fulvum* Ecp2) effector proteins (Stergiopoulos et al. 2012). Some LysM domain-containing proteins formed a monophyletic group within the A-V sub-clade, and their LysM domain is similar to bacterial spore assembly proteins. Almost all other remaining LysM-containing chitinases, which include the C-I, one A-V member (Talma12_7383), and a B-I (Spoth2_113450) protein, share a different recent LysM common ancestor. There is evidence of additional LysM duplications of various ages that have resulted in phylogenetic diversity among LysM domains in the same protein (Supplemental Data 2), however the amino acid alignment is lacking sufficient characters to infer robust relationships among most LysM domains.

Chitin binding domains (CBDs) are dispersed across chitinases in early diverging fungi, Ascomycota and Basidiomycota (Figure 3 and Supplemental Data 2). Zoopagomycota (particularly Basidiobolomycetes) have comparatively large numbers of CBDs, and other early diverging fungi (Mucoromycotina and Kickxellomycotina) have somewhat fewer. CBDs are widely dispersed through the classes and orders of Ascomycota, but in Basidiomycota, they are mostly restricted to Agaricomycetes (Figure 3). The earliest diverging B sub-clade, B-V, has a low percentage of chitinases with CBDs, while there are more of these domains in other sub-clades, particularly the B-III clade (Figure 3). In the AC super clade, most CBDs are found in the C-I and C-II sub-clades as well as in a low percentage of A-V class proteins (Figure 3).

### *H. capsulatum* chitinases

#### Diversification and taxonomic distribution

The *H. capsulatum* genome encodes eight chitinase (Cts) enzymes, which are widely distributed in the GH18 phylogeny (Figure 4 and Supplemental Data 6). Six genomes from diverse *H. capsulatum* strains were used for analysis (*H. capsulatum* G186AR, *H. capsulatum* G217B, *H. capsulatum H143, H. capsulatum* H88, *H. capsulatum* NAM1, and *H. capsulatum* TMU). While the genomes generally contain the same chitinases, there are slight differences in clade distribution*. H. capsulatum* H143 and H88 (two representatives of the African strains) do not contain a C-I chitinase while G217B has two that are 100% identical (likely the result of an assembly error). *H. capsulatum* TMU lacks the otherwise conserved A-IV chitinase, while *H. capsulatum* H143 is also missing the B-I chitinase. We cannot at this time rule out the absence of individual chitinases being due to errors in genome assembly (particularly with the low coverage of H143 at only 3-fold).

**Figure 4.**
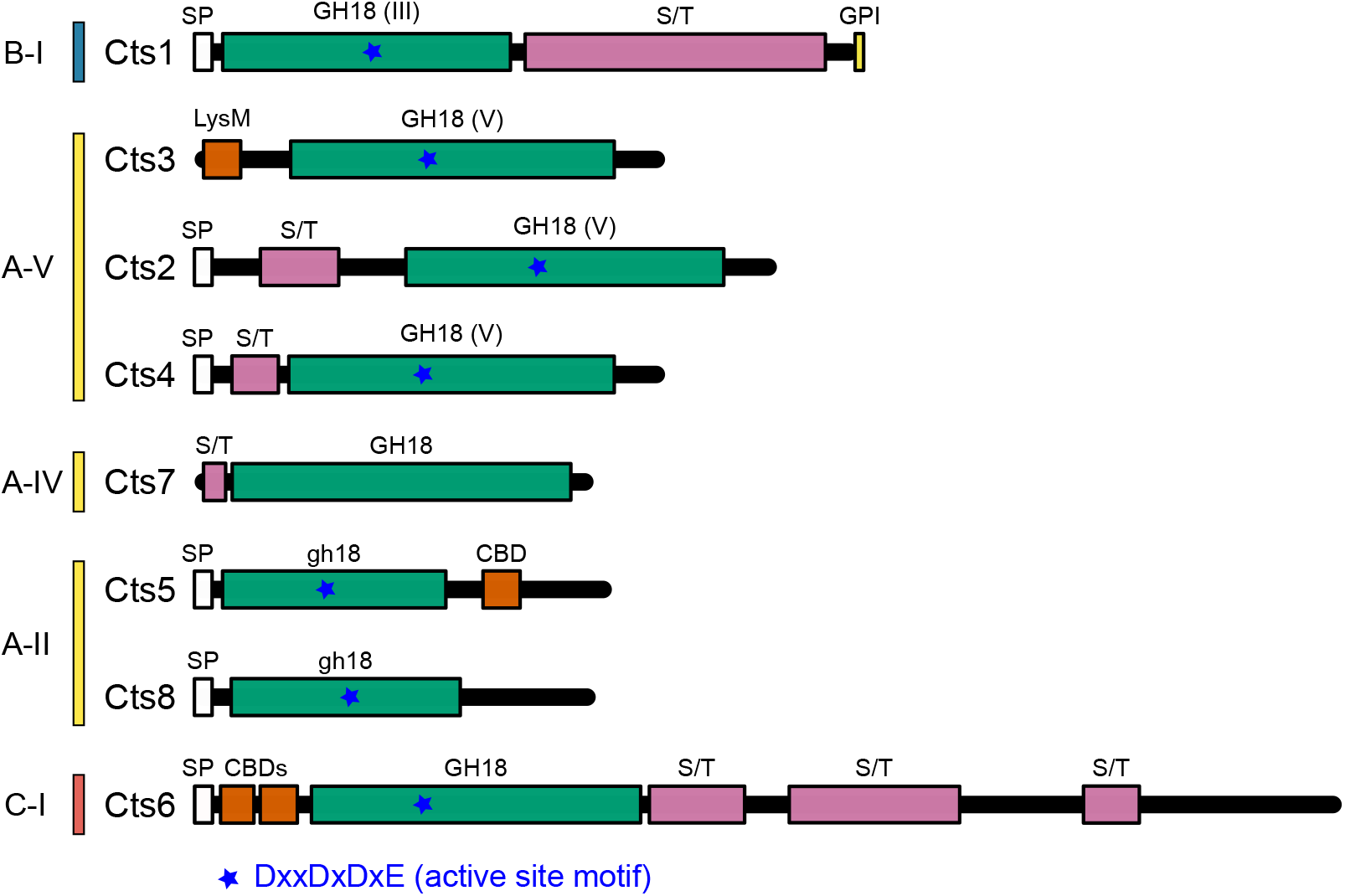
Domain and motif structure of *H. capsulatum* chitinases. The eight chitinase proteins (Cts) encoded in the *H. capsulatum* genome are schematically shown and grouped by sub-clade. Glycosyl hydrolase family 18 (GH18) domains (green) are indicated and noted as class III (GH18(III), class V (GH18(V)), or those with shortened length (gh18). Recognized chitin-binding domains (CBD and LysM) are indicated (orange). N-terminal signal peptides (SP; white) directing secretion of the protein, serine/threonine-rich regions (S/T; pink) as potential sites for O-linked glycosylation, and the motif for glycosylphosphatidylinositol attachment (GPI, yellow) are indicated. Presence and location of residues for the conserved DxxDxDxE motif for chitin hydrolysis is also shown (blue stars). Sub-clade to which each protein belongs is presented along the left.

Cts1 is the only *H. capsulatum* sequence in the B-I sub-clade. Cts6 is the sole C sub-clade protein (C-I) and also has the larger size characteristic of this sub-clade. The remaining *H. capsulatum* chitinases (Cts2, Cts3, Cts4, Cts5, Cts7, and Cts8) are all members of clade A (Figure 4). Cts2, Cts3, and Cts4 are members of the A-V sub-clade, Cts5 and Cts8 both belong to sub-clade A-II, and Cts7 is a member of sub-clade A-IV. Cts4 appears to be a very recent duplication of Cts2 as the closely related fungus *Blastomyces dermatitidis* has only a single Cts protein that is orthologous to both *H. capsulatum’s* Cts2 and Cts4 proteins. *Histoplasma* is the only genus in the Onygenales that contains chitinases in the A-II sub-clade, while other Onygenales tend to contain B-II sub-clade members, which *Histoplasma* lacks. There are no A-III chitinases in *H. capsulatum*, which is consistent with other Onygenales surveyed (Figure 3).

*H. capsulatum* has only a single C-clade chitinase enzyme (C-I) even though the C clade (C-I and C–II) is expanded in multiple other Onygenales members. Among Eurotiomycetes, Onygenales and Eurotiales have similar distributions of chitinases among the sub-clades except as noted above. Chaetothyriales differ in that they contain A-III sub-clade chitinases but lack the B-II and C clade chitinases found in the other Eurotiomycetes (Figure 3).

#### Domain architecture and finer motifs

The architectural diversity of *H. capsulatum* chitinases is consistent with the variability observed across the fungal kingdom. In particular, *H. capsulatum* chitinases look similar to those of other Onygenales. For example, other Onygenales have Cts3-like proteins with matching architecture (i.e, containing a LysM domain but no signal peptide). Most chitinases only have 1-2 CBDs even in C clade proteins. *Microsporum canis* has expanded numbers of CBDs and LysM domains corresponding to their expanded C clade, which is unusual for the Onygenales chitinase architecture. At the individual protein level, domain architecture is variable among the A clade chitinases. Cts2, Cts3, and Cts4 are all in the A-V sub-clade. Of these, Cts2 and Cts4 have similar domain architecture consistent with a recent duplication, with secretion signals at the N-terminus and serine/threonine rich regions (Figure 4), which are common sites for O-linked glycosylation (Loibl and Strahl 2013). Cts3 contrasts with Cts2 and Cts4 by lacking the secretion signal and containing the only LysM domain among *H. capsulatum* chitinases. In sub-clade A-IV, Cts7 lacks a secretion signal and contains a serine/threonine rich region. This is also the only *H. capsulatum* chitinase that lacks conserved aspartate residues predicted to comprise the active site (Hartl et al. 2012). In sub-clade A-II, Cts5 and Cts8 both contain a secretion signal, but only Cts5 contains a recognizable CBD. The C-1 sub-clade chitinase, Cts6, possesses the N-terminal secretion signal, serine/threonine rich regions, and two CBDs. The B clade chitinase in *H. capsulatum*, Cts1 (B-I sub-clade), has a GPI anchor region in addition to a secretion signal suggesting this enzyme is exported and anchored to the cell surface of *H. capsulatum* cells. It also contains an extended serine/threonine rich region common among extracellular proteins. Thus, most *H. capsulatum* chitinases are predicted to be secreted enzymes and active on chitin substrates.

#### Expression of *H. capsulatum* chitinases

To provide insight into the physiological roles of diverse chitinases in *H. capsulatum*, the expression of each was determined under different environmental and nutritional conditions. As a thermally-controlled dimorphic fungus, *H. capsulatum* provides an opportunity to ascribe morphology-specific (yeast- or mycelial-) roles to individual chitinases. Surprisingly, most chitinase-encoding genes (*CTS*) were expressed at constant, but low levels across most conditions including in the presence of exogenous chitin (Figure 5). Only *CTS3* (one of the A-V chitinases) was specifically up-regulated in *H. capsulatum* filamentous cells (on average roughly 11-fold higher in mycelia compared to yeasts; Figure 5) suggestive of mycelia-specific functions. *CTS2* (encoding another A-V-class chitinase) was expressed at higher levels overall suggesting that the Cts2 protein could be important for general growth or the major functional chitinase under the tested conditions. *CTS4* (A-V) and *CTS8* (A-II) were expressed at low but consistent levels. *CTS1* (the only B-I chitinase) showed variable expression. *CTS6* (the C-I chitinase) also showed highly variable levels of expression that somewhat mirrored that of *CTS1* (Figure 5). *CTS7* (the only A-IV chitinase), which lacks the active site D-X-X-D-X-D-X-E residues (Figure 4), showed very low levels of expression. *CTS5* (A-II) expression was not detectable under any condition tested. Interestingly, the presence of exogenous chitin in the media did not consistently induce chitinase gene expression, with the exception of *CTS6* which was induced in the presence of chitin in most conditions except yeast-phase growth on rich media (Figure 5). Regardless, neither *H. capsulatum* yeast nor mycelia were able to grow on chitin as the carbon source of the growth media (data not shown). Thus, with the exception of *CTS3* and *CTS6*, expression studies did not reveal specific environmental conditions for *H. capsulatum* chitinase expression or a yeast-phase specific chitinase.

**Figure 5.**
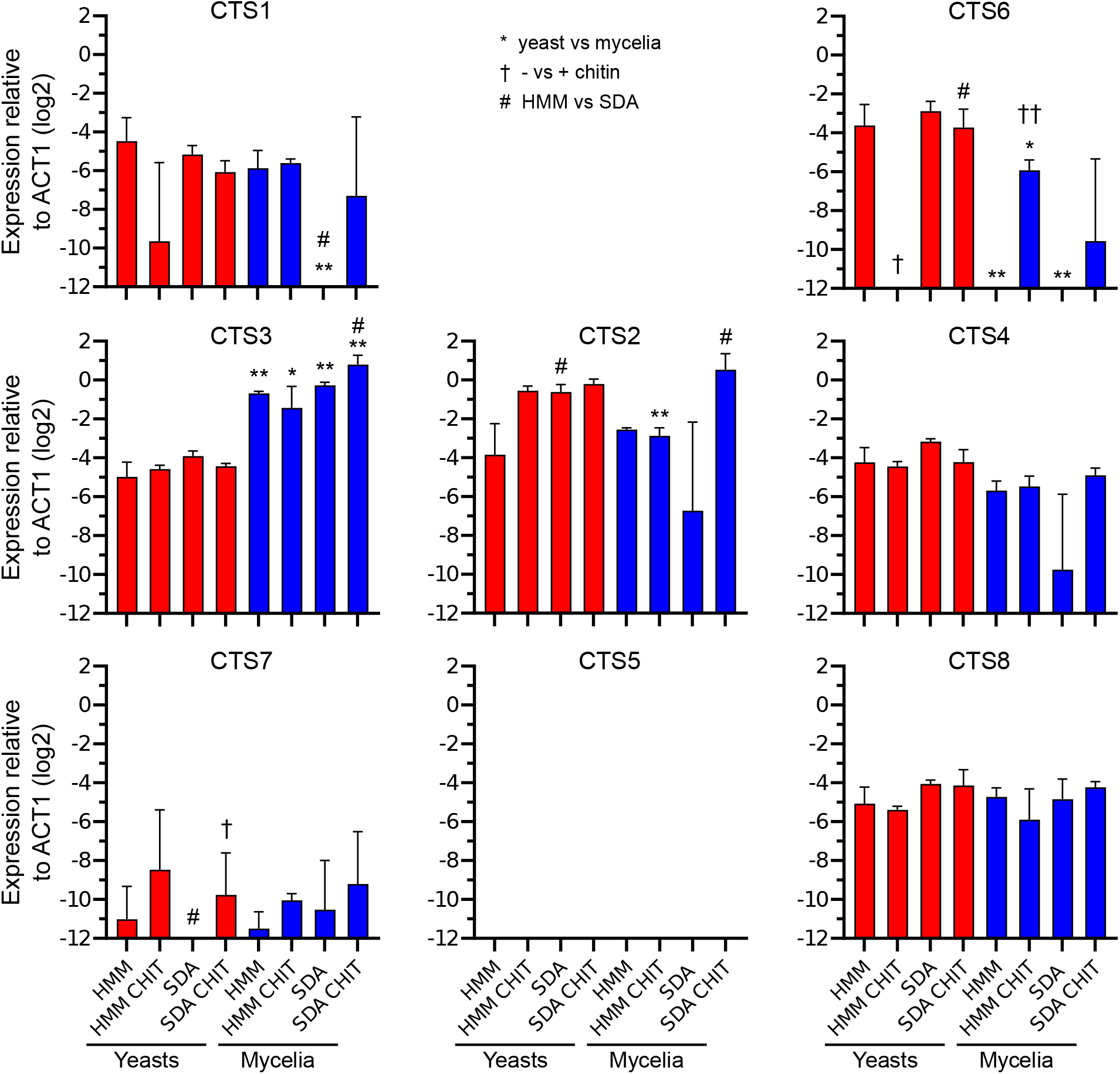
Expression of chitinase-encoding genes by *H. capsulatum* yeasts and mycelia. Expression of each chitinase-encoding gene (*CTS*) by *H. capsulatum* cells was measured using qRT-PCR. Transcription was determined for *H. capsulatum* cells grown on two media, one based on basal cell culture medium (HMM) and one based on glucose and peptone (Sauboraud’s dextrose; SDA). For some tests, media was supplemented with chitin (CHIT). Yeast (red bars) and mycelial (blue bars) phases were maintained by growth at 37°C or 25°C, respectively. Data shows expression of each *CTS* gene relative to the expression of actin (*ACT*) and organized by sub-clades (B, cyan; A, magenta; C, green). Data represent the average ± standard deviation among biological replicates (n=3). Significant differences in expression between yeasts and mycelia (*), absence and presence of chitin (†), or HMM compared to SDA (#) are indicated for corresponding conditions differing by the single parameter. Single symbols indicate P<0.05 and double symbols indicate P<0.01 as determined by pairwise Student’s t-tests.

#### Enzymatic activities of *H. capsulatum* chitinases

To determine if the clades represented by the *H. capsulatum* chitinases (Cts proteins) correspond to different enzyme activities as suggested by previous phylogenetic studies, all eight *H. capsulatum* chitinases were purified and tested for chitin degradation profiles. Three artificial substrate mimics were used to determine the specificity of each purified chitinase enzyme (Figure 6): N-acetyl-glucosaminidase exochitinase activity (liberation of 4-methylumbelliferone (4-MU) from all substrates including sequential hydrolysis of GlcNAc units from the reducing end of oligosaccharides), endochitinase activity (liberation of 4-MU from oligosaccharides with at least 2 GlcNAc units), chitobiosidase activity (hydrolysis of the disaccharide chitobiose from the reducing end thus liberating 4-MU from 4MU-(GlcNAc)2), and chitobiase, which hydrolyzes only the disaccharide chitobiose (e.g., hydrolysis only of the glucosidic bond in 4-MU-GlcNAc). The B-clade chitinase Cts1 exhibited endochitinase activity (Figure 6) consistent with the proposed activity suggested by the more open-channel B-clade enzyme structure (Hartl et al. 2012). The A-V chitinases (Cts2, Cts3, and Cts4) also showed endochitinase activity and chitobiosidase function consistent with hydrolysis of internal glucosidic linkages. Cts4 also had a low level of exochitinase activity similar to N-acetyl-glucosaminidases suggesting neofunctionalization after duplication from Cts2. Of the other A-clade proteins, only Cts8 (an A-II sub-clade protein) exhibited significant chitinase activity, which was consistent with chitobiase activity (Figure 6). The other A-II sub-clade protein, Cts5 had detectable but extremely poor activity on any substrate. Cts7 (sub-clade A-IV), lacked any detectable activity (Figure 6), consistent with the lack of active site residues in the Cts7 protein (Figure 4). Cts6, *H. capsulatum’s* only C-clade protein (C-1 sub-clade) hydrolyzed all substrates consistent with N-acetyl-glucosaminidase exochitinase activity (Figure 6).

**Figure 6.**
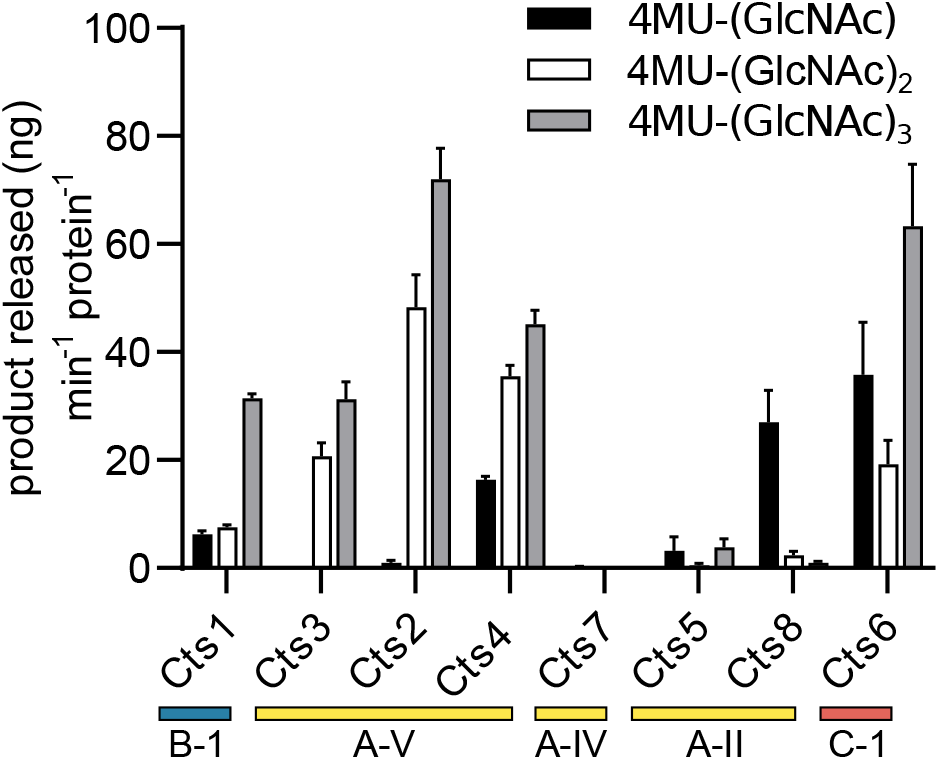
Enzymatic activity of *H. capsulatum* chitinases. The enzymatic activities of purified *H. capsulatum* chitinases were tested using fluorigenic GlcNAc oligomers linked to 4-methylumbelliferone (4MU) by a glycosidic bond. Chitinase activity that hydrolyzes this linkage liberates 4-MU which was detected by fluorescence. Substrates tested were 4MU-(GlcNAc) (black bars), 4MU-(GlcNAc)2 (white bars), 4MU-(GlcNAc)_3_ (gray bars). The exochitinase activity, N-acetyl-glucosaminidase, processively removes GlcNAc saccharides from the non-reducing end resulting in hydrolysis of all substrates. Chitobiosidase activity (release of chitobiose from the non-reducing end) is detected as hydrolysis of 4MU-(GlcNAc)2. Endochitinase activity generates fluorescence from 4MU-(GlcNAc)2 and 4MU-(GlcNAc)3. Chitobiase activity (hydrolysis of the disaccharide chitobiose) generates fluorescence only from the chitobiose mimic 4MU-(GlcNAc). Activity rates were calculated as nanograms of hydrolysis product (4MU) released per minute per nanogram of *H. capsulatum* protein. Data represent the average activity ± standard deviation of replicates (n = 3). GH18 sub-clades to which each Cts protein belong are indicated (A, yellow; B, blue; and C, red).

## DISCUSSION

### Previous chitinase ontologies are largely robust to increased sampling, but the A clade is polyphyletic

Our analyses support the existence of two major classes of chitinases defined by an ancient divergence between B and AC seen in previous phylogenetic analyses. We also recapitulate the previous sub-clades with varying degrees of support (Figure 1 and Table 1). However, we find that there is not a single A clade, because C clade chitinases are more closely related to A-II chitinases. The divergence of the A-IIIs and the close clustering of the A-II sub-clade with the C clade suggest the definition of A and C clades could use revision. Since A clade chitinases are not monophyletic, either C clade chitinases should be subsumed into a larger A clade, or alternately, the A clade chitinases could be split into multiple new clades informed by additional gene architecture and enzymatic activity investigations. Previous work has supported both an independent C clade (Seidl et al. 2005; Karlsson and Stenlid 2008; Alcazar-Fuoli et al. 2011) and a C + A-II clade (Karlsson and Stenlid 2009) that is consistent with our analyses. However, only the present analysis strongly supports the divergence of the A-III clade from the rest of the A sub-clades and the C clade, perhaps because the limited sample of A-III chitinases in previous analyses (< 10 members) did not reflect overall A-III diversity. Our analysis suggests additional fungal chitinases need to be characterized in order to accurately describe functional diversity among chitinases and thereby more accurately inform the designation of clades and sub-clades. Additionally the location of the LysM domain has been used to distinguish the C clade from the A clade (Gruber, Vaaje-Kolstad, et al. 2011; Gruber and Seidl-Seiboth 2012); however, in our analyses, LysM domains are found in the C-II sub-clade as well as some A-V chitinases, including the *H. capsulatum* Cts3 (Figure 4). Furthermore, C-I sub-clade chitinases often lack a detectable LysM domain (e.g., *H. capsulatum’s* Cts6). Thus, the presence of LysM is not a C-clade-defining feature as originally proposed. Additionally, the A clade was generally thought to be lacking in CBDs (Hartl et al. 2012), however CBDs are widespread in the A-II sub-clade (Figure 3), further supporting the need to redefine the A and C clades.

### Species representation within chitinase sub-clades are explained by phylogeny and ecology

The B vs AC split appears to be an early divergence in chitinases, preceding the origin of fungi. For the B clade (GH18 class III) chitinases, previous analyses suggest that B-Vs were the first to diverge after the fungal B chitinases split from bacterial chitinase (Karlsson and Stenlid 2009). Of the AC (Class V) chitinases, sub-clade A-V is the most widespread throughout the taxa, with members found in early diverging fungi, Basidiomycota, and Ascomycota, suggesting it maintains a core function in fungal physiology. In contrast, the A-III sub-clade is greatly reduced in Ascomycota, as compared to early diverging fungi and Basidiomycota, suggesting A-III functions may be conditionally dispensable in Ascomycota. In general, the Pezizomycotina (Ascomycota), particularly Leotiomycetes, Sordariomycetes, Eurotiomycetes, and Dothideomycetes, have chitinases from the greatest diversity of sub-clades. Genomes of early diverging fungi generally contain few chitinases, and these are mostly from the C-II, A-V, and particularly the A-III sub-clades. *Rozella allomycis* is an early diverging species that lacks chitin, but maintains a single A-V chitinase that may play a role in its obligate endoparasitism of chytrid fungi (Jones et al. 2011).

Clearly restricted taxonomic distributions of certain chitinase sub-clades suggest they may be ancestrally orthologous. For example, the B-IV sub-clade is strictly limited to the Saccharomycetales. However, most B-IV characterized functions (Table 2) relate to fungal cell division and morphology (Colussi, Specht, and Taron 2005; Hurtado-Guerrero and van Aalten 2007; Kuranda and Robbins 1991; Selvaggini et al. 2004). As cell division is fundamental, it would be unusual that this function itself is performed only by chitinases in Saccharomycetales. The constitutive expression of *H. capsulatum’s* B-clade chitinase, and the GPI-attachment motif suggestive of anchoring at the cell surface, supports a generalized function for B clade chitinases in fungal cell division and growth. The B-III sub-clade may represent the Basidiomycota-specific B chitinases, but as delimited it also includes Mucorales chitinases, which may reflect uncertainty in deep branching order, or loss of an Ascomycota paralog. B-I and B-II chitinases are limited to the Pezizomycotina and may represent a taxon-specific radiation of B chitinases.

**Table 2.**
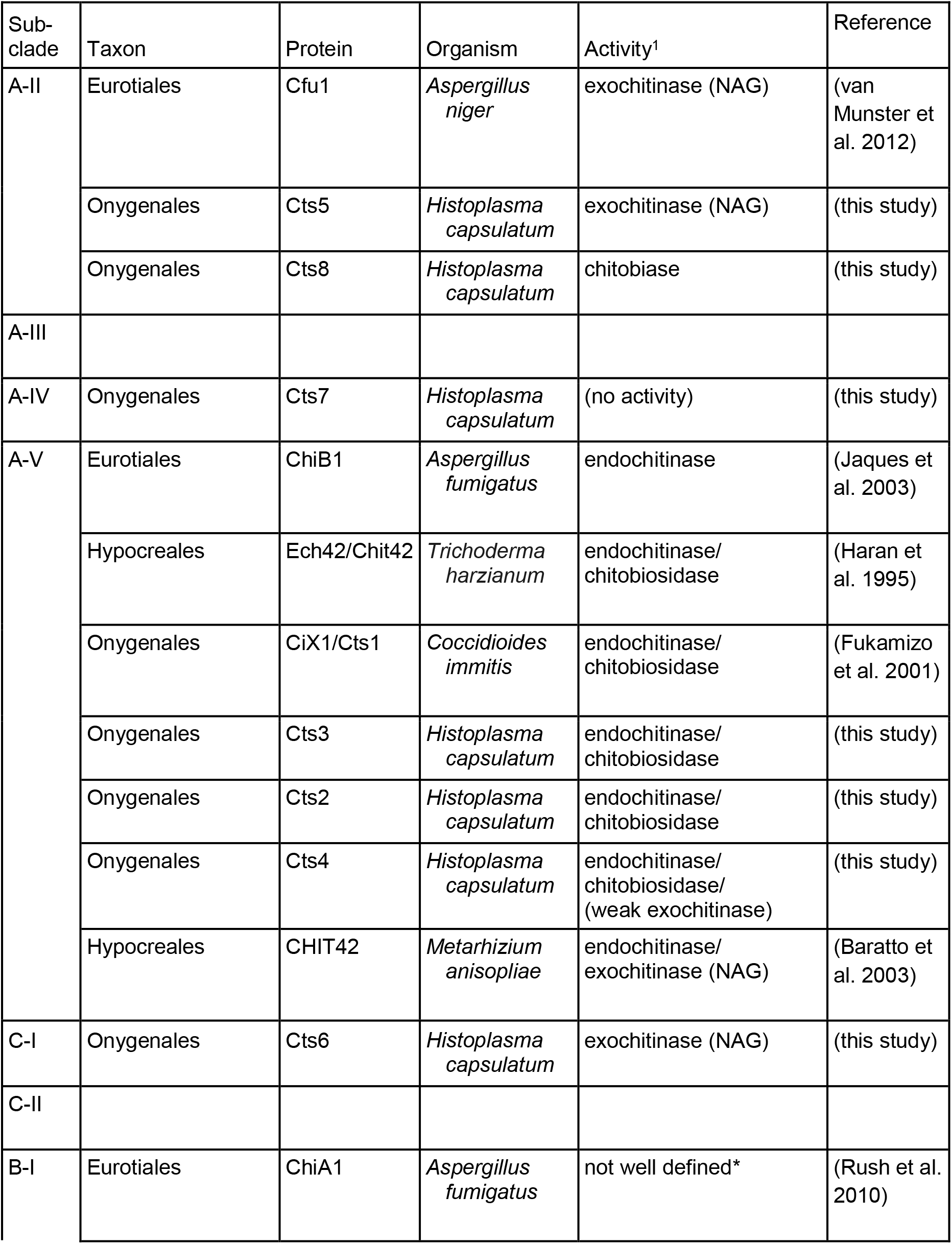
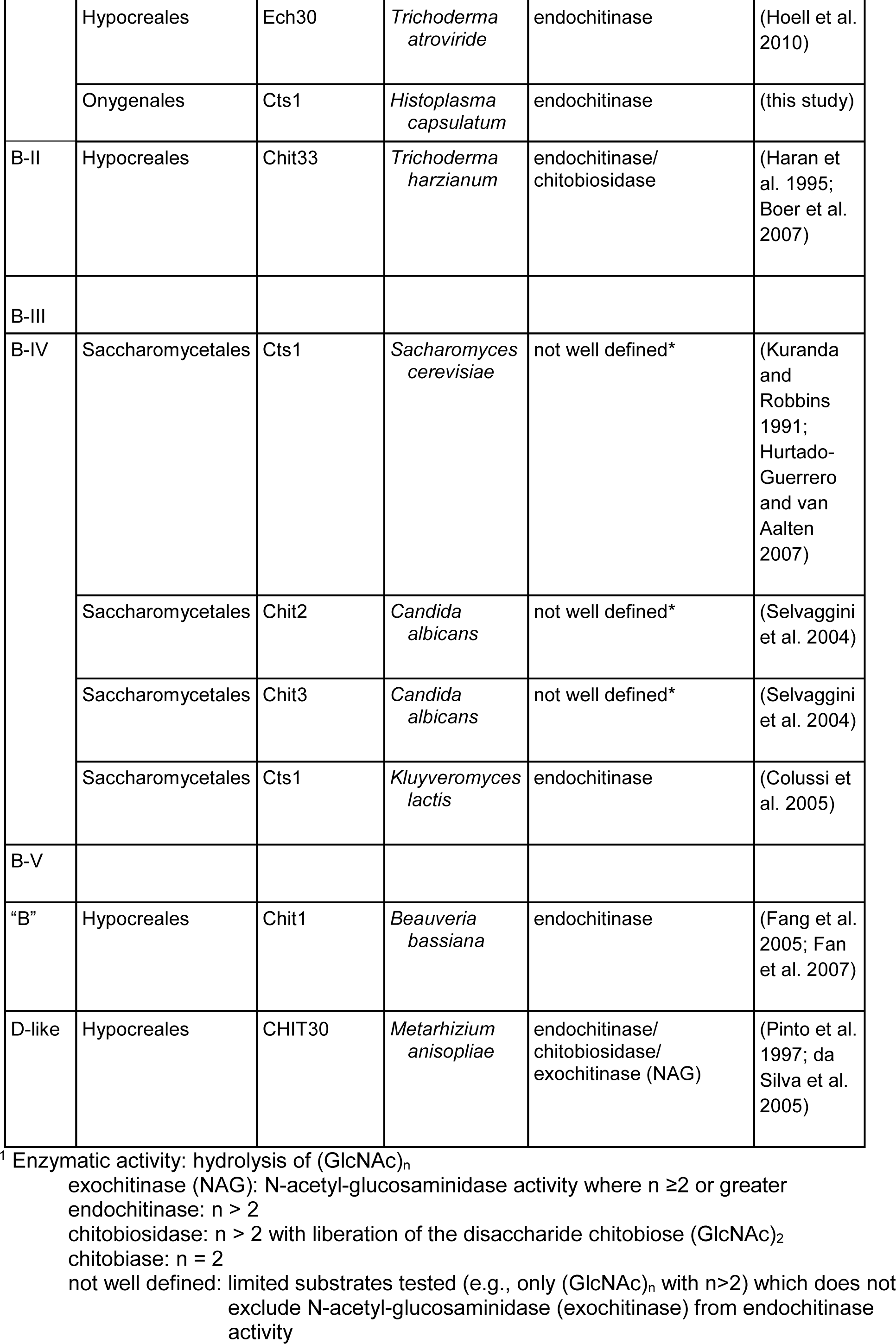
Enzymatically Characterized Proteins.

The fungal ecological diversity represented among chitinase clades is complex, precluding simple ecological explanations for differences among chitinase repertoires; however the variability in the number of chitinases in species is consistent with some expectations. For example, the genomes with the highest number of chitinases include fungi with predominantly insect and fungus-derived nutrition like Basidiobolus (Tabima et al. 2020) Cordyceps, and Trichoderma (Supplemental Data 7), but also wood-decay fungi like Ganoderma and Auricularia, which may use additional chitinases in competition or secondary decomposition. In contrast we see the fewest chitinases (≤1 per sequenced genome) in fungi that are sequestered from the open environment or obligately associated with plants or animals. For example, few chitinases are produced by Rozella (endoparasites), Neocallimastigomycota (obligate gut rumen symbionts), Glomerales (arbuscular mycorrhizae), and Malassezia (skin commensals). We identified no GH18 chitinases in the obligate intracellular animal parasites Microsporidia, and the GH18 function may have been subsumed by their recently described GH19 chitinases (Han et al. 2016).

Although it was not feasible to perform a comprehensive analysis of individual molecular evolution events among fungal chitinases in this dataset, there is evidence for a significant role of HGT in their diversification, and these events suggest ecological roles for the transferred genes (Supplemental Figure 1 and Supplemental Data 5). For example, we observed multiple HGT events and gene family expansions among insect-associated fungi. As insect exoskeletons are a major source of chitin (Tharanathan and Kittur 2003), the acquisition and expansion of chitinases might facilitate greater chitinase production or enable degradation of diverse forms of arthropod chitin under diverse environmental conditions or pathogen relationships (Karlsson and Stenlid 2009). LysM domain-containing chitinases are distributed among plant pathogenic, insect parasitic and saprotrophic fungi.

LsyM domains have been shown to bind chitin although carbohydrate-binding specificities are not well-described and their role in most fungi is not well understood (Gruber, Vaaje-Kolstad, et al. 2011). In our analysis, the LysM domain, although largely corresponding to the C-II clade, is not a clade-defining characteristic, making prediction of the functional role of such chitinases difficult (Figure 3). Some LysM-containing proteins have also been shown to bind fungal chitin, contributing to fungal evasion of host plant immunity (Bolton et al. 2008; de Jonge and Thomma 2009). One chitinase in the single Chytrid genome analyzed, *Spizellomyces punctatus*, which may be a decomposer of arbuscular mycorrhizal fungi (Paulitz and Menge 1984) contains a large number of LysM domains. However, although there are many plant and animal pathogens with LysM-containing chitinases, there does not appear to be a strong ecological association (Stergiopoulos et al. 2012).

### Evolution of the *H. capsulatum* chitinases indicates a degree of differentiation and expansion that is reflected in fungal chitinases in general

Placing *H. capsulatum* in context of the other Onygenales analyzed, *H. capsulatum* was the only genus that contained A-II sub-clade members. One of the most interesting features of *H. capsulatum* chitinase evolution is the presence of two chitinases in the A-II sub-clade. A-II sub-clade members are not widespread in fungal taxonomy; they are almost exclusively found in the Leotiomyceta (sub-division of the filamentous Pezizomycotina). A-II chitinases are found in Eurotiomycetes, Sordariomycetes, Dothideomycetes, and Leotiomycetes. In the Eurotiomycetes, Eurotiales, Onygenales, and Chaetothyriales contain A-II sub-clade members. However, *H. capsulatum* is the only Onygenales to contain this sub-clade. That indicates these chitinases are ancestral to the Leotiomyceta, however they have been largely lost in the Onygenales with the exception of *H. capsulatum*. Therefore, these may have a specific function that is necessary for *H. capsulatum* biology that is lacking in other Onygenales.

The B-clade chitinase in *H. capsulatum* (Cts1) belongs to the B-I clade. The other Onygenales tend to contain B-II sub-clade members that *H. capsulatum* lacks. In addition, the C clade chitinases are expanded in multiple other Onygenales members but is restricted to one chitinase in *H. capsulatum* (Cts6). This is most obvious in *Microsporum canis* with 9 C-I and 4 C-II members. However, upon closer examination, only 3 of each sub-clade are predicted to be active (contain the active site residues) indicating that this larger clade representation may not be as functionally extreme as otherwise indicated.

In terms of other members of Eurotiomycetes, the Onygenales and Eurotiales show matching patterns of chitinases distribution in the sub-clades. Chaetothyriales differ in that they contain A-III sub-clade chitinases members but are lacking B-II or C clade members seen in the other Eurotiomycetes. When comparing the Eurotiomycetes to other Pezizomycotina, Saccharomycetes is rather divergent. It is the only one to contain a B-IV domain, and is the only order from the Pezizomycotina to be missing a B-V or C-I sub-clade chitinase from the Pezizomycotina. Saccharomycetes are limited to the B-IV, C-II, A-IV, and A-V sub-clades, which is less diversity seen in other Pezizomycotina. Sordariomycetes and Eurotiomycetes had identical patterns of sub-clade distribution, missing the A-III proteins. Leotiomyceta members (excluding Xylonomycetes) have A-II, B-I, and C-II sub-clade proteins. The A-IV clade is missing in Orbiliomycetes while C-I is missing in Xylonomycetes and Pezizomycetes. The A-V sub-clade is conserved in all Pezizomycotina.

### *H. capsulatum* chitinase expression and functional differences suggest complementary functional roles

*H. capsulatum* expression data allowed for an initial look at how multiple chitinases could potentially play different roles. Cts1 is secreted and contains a GPI-attachment motif indicating that it is likely anchored at the fungal cell wall for cell wall remodeling or modification during cell division and growth. Cts1 had a low, but generally constant level of expression in all growth conditions consistent with an essential function in fungal growth. As transcriptional data was not collected on synchronized stages of division or morphology, it is possible the low expression is a result of a mix of cells at different stages of replication with high and low expression. The finding that Cts1 is an endochitinase active only on larger oligosaccharides (Figure 6) rather than a processive exochitinase fits with a putative role in remodeling, but not complete dismantling, of chitin in the fungal cell wall.

Cts3 was the only chitinase that was induced in the mycelial phase of *Histoplasma*, supporting a functional role in the mycelial state of this organism, which is an environmental saprotroph in contrast to the yeast phase that is parasitic for animals. As Cts3 lacks a secretion signal, this may suggest that Cts3 plays an intracellular role in cell division, septa formation, or in the preparation of the cell wall for formation of mycelial-specific structures (e.g., conidiophores). The chitobiosidase activity of Cts3 suggests the existing cell wall must be extensively altered for such structures. Chitinases are generally expanded in fungi characterized by filamentous growth and are few in yeast-form fungi like *S. cerevisiae*, supporting the hypothesis that hyphal growth or reproductive morphologies requires multiple different chitinases (Karlsson and Stenlid 2008). As Cts3 is the only *H. capsulatum* chitinase with a LysM domain, this may further indicate that Cts3 activity is directed at “self” chitin (e.g., during formation of hyphal wall structures) similar to the role of LysM-proteins in binding fungal pathogen chitin to prevent plant immune responses. *H. capsulatum* dimorphism is strictly regulated (Edwards et al. 2013), primarily controlled by the Ryp transcription factors (Shen and Rappleye 2017; Sil 2019) while other common fungal transcription factors, such as the APSES family, appear to have very limited function in dimorphism (Longo et al. 2018). Therefore, any chitinases necessary for hyphal growth or mycelial structures should be closely linked to transcriptional regulation of the mycelial state.

*CTS2*, which encodes another A-V sub-clade chitinase, has consistently higher expression among all *H. capsulatum* chitinase genes. This suggests that under the conditions tested Cts2 may be the general functional chitinase while the other related chitinase, Cts4, is either simply less used or used under specific, unknown conditions. Other *H. capsulatum* chitinases show low or highly variable expression among diverse conditions precluding specific hypotheses as to their biological roles based on gene expression.

As *H. capsulatum* is the only Onygenales member with A-II chitinases, the secreted Cts5 and Cts8 chitinases are of particular interest. Cts8 hydrolysis activity is consistent with that of a chitobiase, an enzyme that specifically hydrolyzes the disaccharide chitobiose which is produced by the activity of chitobiosidases. This may indicate that the primary role of Cts8 is to further hydrolyze disaccharides in the extracellular environment (such as those produced by other chitinases) into GlcNAc monomers. We hypothesize that Cts8 thus plays a nutritional role by liberating hexosamines from other chitin hydrolysis products. Alternatively, generation of GlcNAc monomers from environmental chitin degradation products (i.e., chitobiose) may serve a signaling function since adoption of the mycelial state is potentiated by free GlcNAc but not glucosamine (Gilmore et al. 2013). The other A-II *Histoplasma* chitinase, Cts5, has very minimal activity, which along with the lack of any detectable expression suggests that this chitinase may no longer have a functional role.

*CTS7* expression is very low to undetectable, and Cts7 lacks any chitinase activity suggesting that it also may no longer serve a functional role in *H. capsulatum* biology. Consistent with this, Cts7 also lacks a signal sequence for secretion unlike the majority of the other chitinases. However, at this time a non-chitin-hydrolysis role for Cts7 cannot be ruled out.

Cts6 is the most unique of *H. capsulatum’s* chitinases having multiple domains and an increased size. Cts6 is the only C-clade chitinase produced by *H. capsulatum*, and the expression pattern of *CTS6* suggests that its transcription increases in the presence of exogenous chitin. Together these characteristics suggest that Cts6 may be suited for hydrolysis of diverse chitin substrates in the environment. Consistent with a role in the degradation of environmental chitin, Cts6 is secreted and has N-acetyl-glucosaminidase activity (Figure 6), an exochitinase function which would enable it to processively hydrolyze chitin polysaccharides.

### *H. capsulatum* chitinase specificity demonstrates the difficulty in using phylogeny for activity prediction

Determination of the chitinase activities of the eight *H. capsulatum* chitinases highlights the limitations of previous phylogenetic analyses to predict enzymatic activities. Few fungal chitinases have been enzymatically characterized (Table 2) and the characterization is sometimes not complete. The multiple A-V *H. capsulatum* chitinases illustrate how functional variation can be present even within sub-clades, such as the exochitinase activity of Cts4 in addition to the endochitinase and chitobiosidase activity of its closest paralog Cts2. When compared to some other characterized A-V clade members we see a similar pattern of multiple and complex activities (Table 2). The demonstrated endochitinase activity with Cts2 and Cts3 is inconsistent with the previous prediction that A clade members are characterized by exochitinase activity (Duo-Chuan 2006; Seidl 2008; Hartl et al. 2012). However, this sub-clade also shows a strong presence of chitobiosidase activity (Table 2). The A sub-clades might have functionally diverged, which may be reflected in the potential polyphyletic phylogeny of the A clade chitinases. For example, the characterized members of the A-II sub-clade have varied activities, not the previously suggested restriction to exochitinase activity (Table 2). In addition, the A-III sub-clade is lacking in enzymatically-characterized members, and the A-IV only has one characterized member (Cts7 from this study), which does not have any enzymatic activity at all. As the A sub-clades are more variable in enzymatic activity than expected and the A-III subclade is so divergent from the other AC sub-clades, these chitinases need further study to uncover how the A sub-clades have diverged and how that might influence their activities. This study is the first report of the enzymatic activity of a C clade member (Table 2). The hydrolysis profile of *H. capsulatum* Cts6 is consistent with an N-acetyl-glucosaminidase as the successive hydrolysis of GlcNAc from oligosaccharides enables this enzyme to hydrolyze all substrates tested (Figure 6).

In contrast to the AC clade, the B clade functional data is much more consistent with predictions, with characterized members of multiple clades having endochitinase or chitobiosidase activity (Duo-Chuan 2006; Seidl 2008; Hartl et al. 2012). However, sub-clades B-III and B-V have not been extensively investigated to determine the universality of the prediction for B clade enzymes (Table 2). In addition, some of these functionally characterized enzymes have not been extensively studied for substrate preference and thus a strict categorization is premature.

The large increase in available fungal genomes is improving our understanding of the evolutionary relationships among fungal chitinases. As demonstrated in this study, combination of phylogenetics with empirical characterizations, including determination of enzymatic activities, further refines the predictive power of fungal chitinase phylogenetic classifications. Using this fungal chitinase phylogenetic framework, future studies to define the functional roles of enzymes using the increased feasibility of molecular genetic tools for fungi (Wang et al. 2017; Raschmanová et al. 2018; Song et al. 2019) will connect evolution of chitinases to their individual and specific roles in fungal biology.

## MATERIALS AND METHODS

### Phylogenetic Analyses

Putative chitinases were retrieved from 373 published fungal genomes, including 6 *Histoplasma* genomes (Supplemental Data 1) obtained from the Joint Genome Institute (JGI) website and NCBI. Proteomes were independently mined for glycosyl hydrolase 18 (GH18) containing proteins (El-Gebali et al. 2019) using hidden Markov models (HMM search) (Eddy 2009). A CDD search was then performed on proteins containing GH18 domains to identify additional carbohydrate binding domains (Marchler-Bauer et al. 2017) and LysM domains. Alignments of the GH18 domains and LysM domains were performed by HMMalign which uses a Viterbi algorithm to align each sequence to the given HMM (Eddy 2009). Sequences were removed if they were missing alignment between positions 87-238. Poorly aligned characters were removed using trimAl (v. 1.4) (Capella-Gutiérrez et al. 2009) with a gap threshold of 0.01.

Phylogenetic analysis of the GH18 domains was performed using maximum-likelihood methods using VT+F+G4 (GH18) or WAG+R5 (LysM) model (automatically determined) in IQ-TREE (Nguyen et al. 2015). This tree was used to determine the sequences in the AC clades and the B clade. These sequences were then split into separate phylogenetic analyses again using maximum-likelihood methods (model WAG+G4 for AC tree and PMB+F+I+G4 for 4B tree as automatically determined) implemented in IQ-TREE (Nguyen et al. 2015). Statistical support for all IQ-TREEs was assessed by Ultrafast bootstrap analysis using 1,000 replicates (Minh et al. 2013). An ultrafast bootstrap value of ≥ 95 was considered a strongly supported branch. Parameters of additional phylogenetic analyses in IQ-TREE to assess support for HGT events, including topology tests using the Approximately Unbiased test (Shimodaira 2002) implemented in IQ-TREE are in Supplemental Data 5. In addition, maximum-likelihood methods were implemented in RAxML v.(SSE3) (Stamatakis 2014), 100 alternative runs on 100 distinct trees, bootstrapping (100) and best scoring ML in 1 run and GAMMA models autodetermined (WAG for AC and PMB for B). Statistical support for RAxML trees was assessed by rapid bootstrapping, where nodes receiving ≥60 percent of bootstraps were considered supported.

### Chitinase gene expression

*H. capsulatum CTS* gene transcription was determined by quantitative RT-PCR (qRT-PCR). The WU15 strain of *H. capsulatum*, a uracil auxotroph of the North America 2 clade was grown on HMM (Worsham and Goldman 1988) or Sabouraud’s Dextrose Agar (SDA) media supplemented with 100 μg/mL uracil and solidified with 0.6% agarose. For tests of chitin induction of *CTS* gene transcription, colloidal chitin (prepared from shrimp chitin (Rodriguez-Kabana et al. 1983)) was added to media (1.2% final concentration). Media were inoculated with *H. capsulatum* cells and incubated at 25 degrees (mycelial culture) for 4 weeks or 37 degrees (yeasts culture) for 5-7 days. *H. capsulatum* cells were scraped from the solid media and collected by centrifugation (5 minutes at 2000 x g). RNA was isolated from fungal cells by mechanical disruption with 0.5 mm glass beads, extraction with RiboZol (Amresco), and purification using an affinity column (Direct-zol RNA MiniPrep Plus; Research Products International). Following DNA removal with DNase Invitrogen, RNA was reversed transcribed (Maxima reverse transcriptase; Thermo Scientific) primed with random pentadecamers. Quantitative PCR was carried out using *CTS* gene specific primer pairs (Supplemental Data 8) with SYBR green-based visualization of product amplification (Bioline). Changes in *CTS* gene transcript levels relative to Actin (*ACT1*) and Ribosomal Protein S15 (*RPS15*) were determined using the ΔΔCt method (Schmittgen and Livak 2008) after normalization of cycle thresholds to *ACT1* and *RPS15* mRNA levels. RNAs were prepared from 3 biological replicates for each media/condition. Significant differences in expression (P<0.05) were determined by Student’s t-test between paired samples differing by a single parameter. Samples with transcripts below the detectable limit were set to −12.00 ΔΔCt for analysis purposes.

### Purification of *H. capsulatum* chitinases

Chitinases were purified from transgenic *H. capsulatum* yeasts overexpressing each protein. Chitinase-encoding genes were amplifed by PCR from wild-type *H. capsulatum* G217B genomic DNA and cloned into the *H. capsulatum* expression vector pAG38 placing the gene under transcriptional control of the *H2B* constitutive promoter and fusing the chitinase to a C-terminal hexahistidine tag. For Cts1, sequences encoding the putative GPI-attachment site (nucleotides 2141-2215) were removed to permit recovery of soluble protein. Overexpression plasmids were transformed into *H. capsulatum* WU15 yeasts by *Agrobacterium-mediated* transformation (Zemska and Rappleye 2012) and transformants selected by uracil prototrophy. Transformants were screened by immunoblotting of culture filtrates and cellular lysates for the hexahistidine tag (GnScrpt antibody A00186 to 6xhistidine). Chitinase-expressing transformants were grown in liquid HMM until stationary phase and the yeasts separated from the culture supernatant by centrifugation (5 minutes at 2000 x g). For Cts1, Cts2, Cts4, Cts5, Cts6, and Cts8, culture filtrates were prepared by filtration of the supernatant (0.45 um pore; Millipore), concentrated 100-fold by ultrafiltration (10 kDa MWCO; Whatman), and the proteins exchanged into phosphate-buffered saline (PBS). For Cts3 and Cts7, which lack secretion signals, lysates were prepared from the yeast cells by suspension of yeasts in PBS and mechanical breakage with 0.5 um diameter glass beads. Debris was removed from the cellular lysate by centifiguation (10 minutes at 12000 x g). Hexahistidine-tagged chitinase proteins were purified from the concentrated culture filtrates or cellular lysates by metal affinity chromatography (HisPur Co^2+^ Resin, Thermo Fisher Scientific) and the chitinase-containing elution fractions exchanged into PBS by ultrafiltration. Resultant protein concentrations were determined using a Bradford assay (Sigma-Aldrich) and purity of the protein preparation determined by SDS-PAGE followed by silver staining. Purified proteins were stored at −20 °C in 50% glycerol.

### Chitinase activity and specificity determination

Chitinase enzymatic activities were determined via a fluorimetric chitinase assay kit (Sigma-Aldrich-CS1030). Three artificial substrate mimics, 4-Methylumbelliferyl N-acetyl-β-D-glucosaminide (4MU-(GlcNAc)), 4-Methylumbelliferyl N,N’-diacetyl-β-D-chitobioside (4MU-(GlcNAc)2), and 4-Methylumbelliferyl β-D-N,N’,N’’-triacetylchitotriose (4MU-(GlcNAc)3) were used to determine the activity of each chitinase. Hydrolysis of each substrate releases 4-methylbelliferone, the fluorescence of which was measured using a plate reader (360 nm excitation and 450 nm emission; BioTek Synergy 2). For chitinase activity reactions, varying amounts of purified chitinases were added to 0.5mg/mL substrate in reaction buffer (see kit protocol) and incubated at 37°C. Reactions were monitored by endpoint fluorescence at 60 minutes. Data is reported as nanograms of 4-methylumbelliferone released per minute per nanogram of purified chitinase. The equation of the best-fit line to the data was computed and compared to a standard curve of 4-methylumbelliferone. Activity assays were performed in triplicate.

## ACKNOWLEDGEMENTS AND FUNDING

We thank Stephanie Ray and Qian Shen for technical reviewing and editing of the manuscript. This work was supported in part by grant AI117122 to C.A.R. from the National Institutes of Health and grant DEB-1638999 to J.C.S. from the National Science Foundation. K.D.G was supported in part through a predoctoral fellowship sponsored by the NIH/NIAID award T32, 5T32AI112542-05 (MPI), a NRSA training grant administered by the Department of Microbial Infection and Immunity at Ohio State University and a postdoctoral fellowship sponsored by the NIH award T32, 5T32HL007749-27 at the University of Michigan. The funders had no role in study design, data collection and interpretation, or the submission of the work for publication.

---

**Supplemental Figure 1. Detection of potential horizontal gene transfer occurrances**

Maximum Likelihood phylogenies (IQ-tree) to infer origins of potential HGT chitinases (red text). (**A**) The A clade of chitinases from arthropod pathogens in Hypocreales occupy an uncertain position among diverse bacteria. (**B**) The A clade of mostly Hypocreales chitinases are placed within Streptomyces (Actinobacteria). (**C**) Two clades of fungal chitinases occupy separate parts of the phylogeny of D-like chitinases from Actinobacteria. (**D**) The A clade of Kickxellales chitinases also contains sequences from multiple hypocrealean insect pathogens. (**E**) The B-V chitinase from *Basidiobolus meristosporus* (Zoopagomycota) is placed in Agaricomycetes (Basidiomycota). (**F**) The C-II chitinase in *Panaeolus cyanescens* (Agaricales) is sister to a chitinase in *Uncinocarpus reesei* (Onygenales). Support values indicate percentage of rapid bootstraps.

**Supplemental Figure 2. Phylogeny of LysM domains**

Maximum Likelihood phylogeny (IQ-tree) of LysM domains extracted from all fungal chitinases. Support values indicate percentage of rapid bootstraps.

## Notes

### Competing Interest Statement

The authors have declared no competing interest.

### Summary of Updates

Figure 3 and Supplemental data updated

