## Supplemental Figures for "Diversification of fungal chitinases and their functional differentiation in *Histoplasma capsulatum*"

**Supplemental Figure 1. Detection of potential horizontal gene transfer occurrances**
Maximum Likelihood phylogenies (IQ-tree) to infer origins of potential HGT chitinases (red text). (**A**) The A clade of chitinases from arthropod pathogens in Hypocreales occupy an uncertain position among diverse bacteria. (**B**) The A clade of mostly Hypocreales chitinases are placed within Streptomyces (Actinobacteria). (**C**) Two clades of fungal chitinases occupy separate parts of the phylogeny of D-like chitinases from Actinobacteria. (**D**) The A clade of Kickxellales chitinases also contains sequences from multiple hypocrealean insect pathogens. (**E**) The B-V chitinase from Basidiobolus meristosporus (Zoopagomycota) is placed in Agaricomycetes (Basidiomycota). (**F**) The C-II chitinase in Panaeolus cyanescens (Agaricales) is sister to a chitinase in Uncinocarpus reesei (Onygenales). Support values indicate percentage of rapid bootstraps.

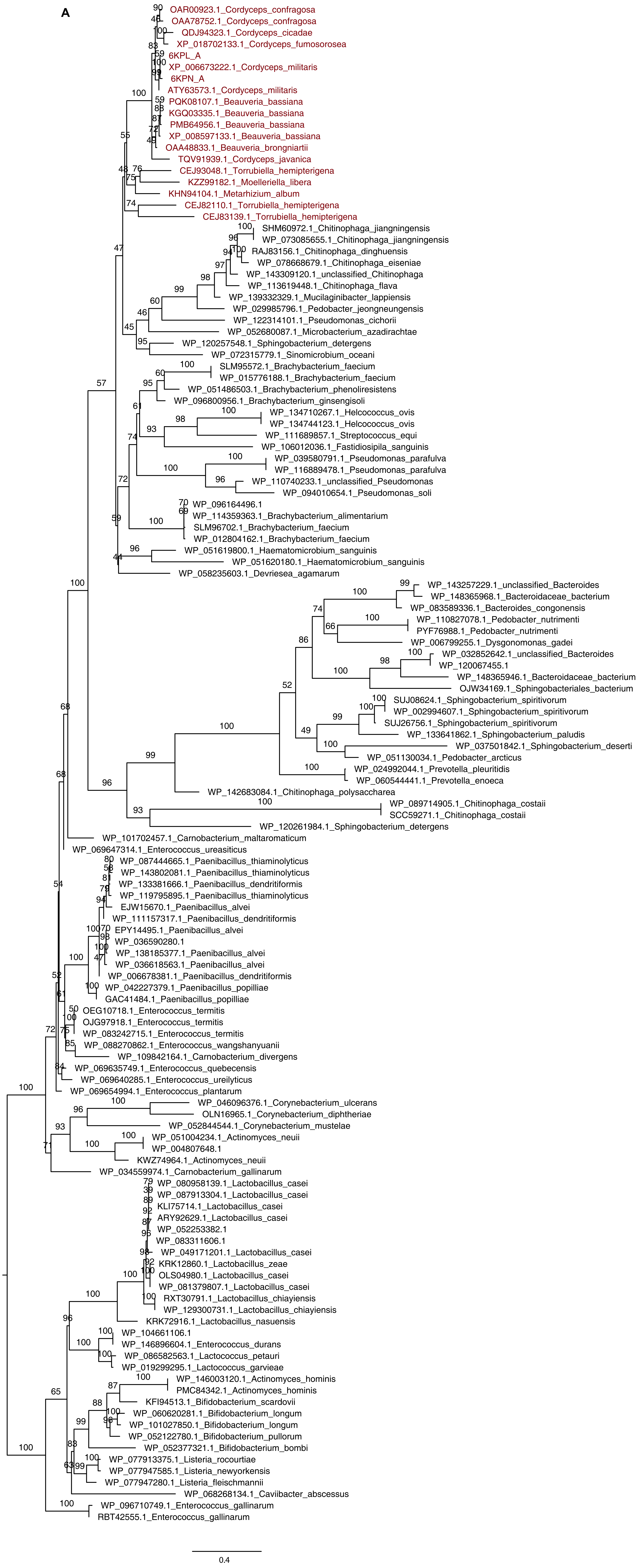

B

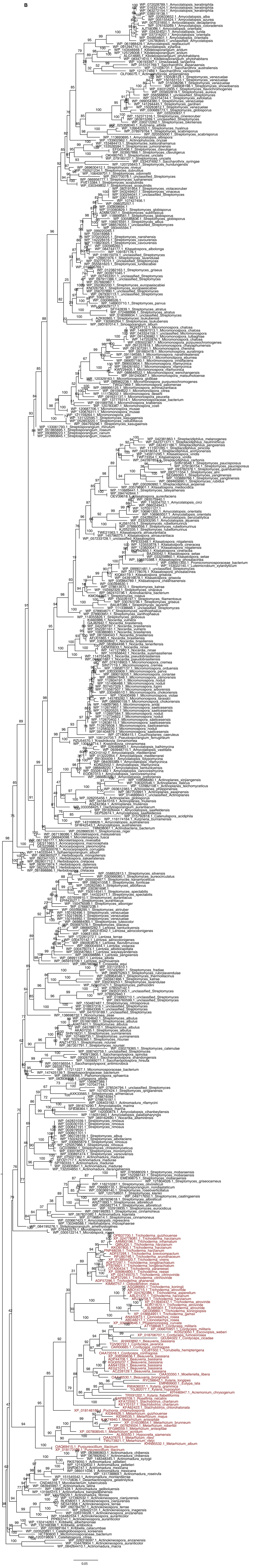

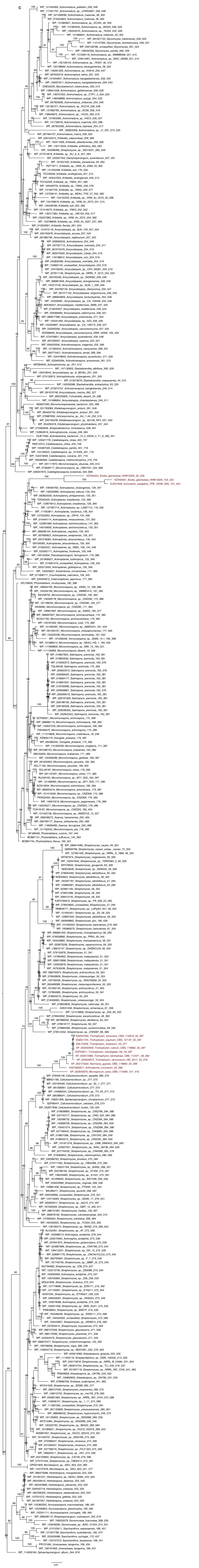

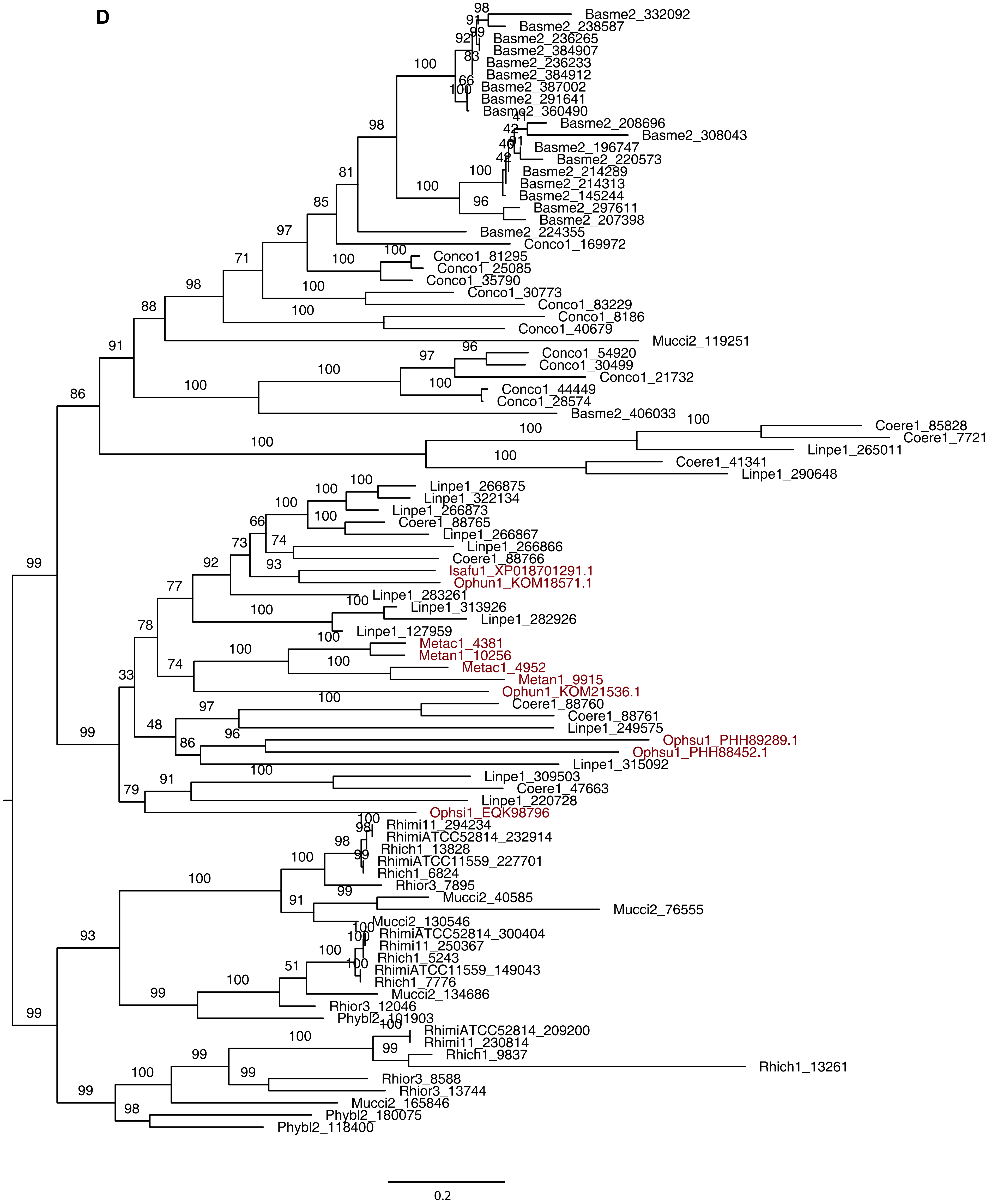
D
Basme2_332092
Basme2_238587
Basme2_236265
Basme2_384907
Basme2_236233
Basme2_384912
Basme2_387002
Basme2_291641
Basme2_360490
Basme2_208696
Basme2_308043
Basme2_196747
Basme2_220573
Basme2_214289
Basme2_214313
Basme2_145244
Basme2_297611
Basme2_207398
Basme2_224355
Conco1_169972
Conco1_81295
Conco1_25085
Conco1_35790
Conco1_30773
Conco1_83229
Conco1_8186
Conco1_40679
Mucci2_119251
Conco1_54920
Conco1_30499
Conco1_21732
Conco1_44449
Conco1_28574
Basme2_406033
Coere1_85828
Coere1_7721
Linpe1_265011
Coere1_41341
Linpe1_290648
Linpe1_266875
Linpe1_322134
Linpe1_266873
Coere1_88765
Linpe1_266867
Linpe1_266866
Coere1_88766
Isafu1_XP018701291.1
Ophun1_KOM18571.1
Linpe1_283261
Linpe1_313926
Linpe1_282926
Linpe1_127959
Metac1_4381
Metan1_10256
Metac1_4952
Metan1_9915
Ophun1_KOM21536.1
Coere1_88760
Coere1_88761
Linpe1_249575
Ophsu1_PHH89289.1
Ophsu1_PHH88452.1
Linpe1_315092
Linpe1_309503
Coere1_47663
Linpe1_220728
Ophsi1_EQK98796
Rhimi11_294234
RhimiATCC52814_232914
Rhich1_13828
RhimiATCC11559_227701
Rhich1_6824
Rhior3_7895
Mucci2_40585
Mucci2_76555
Mucci2_130546
RhimiATCC52814_300404
Rhimi11_250367
Rhich1_5243
RhimiATCC11559_149043
Rhich1_7776
Mucci2_134686
Rhior3_12046
Phybl2_101903
RhimiATCC52814_209200
Rhimi11_230814
Rhich1_9837
Rhich1_13261
Rhior3_8588
Rhior3_13744
Mucci2_165846
Phybl2_180075
Phybl2_118400
0.2

**E**

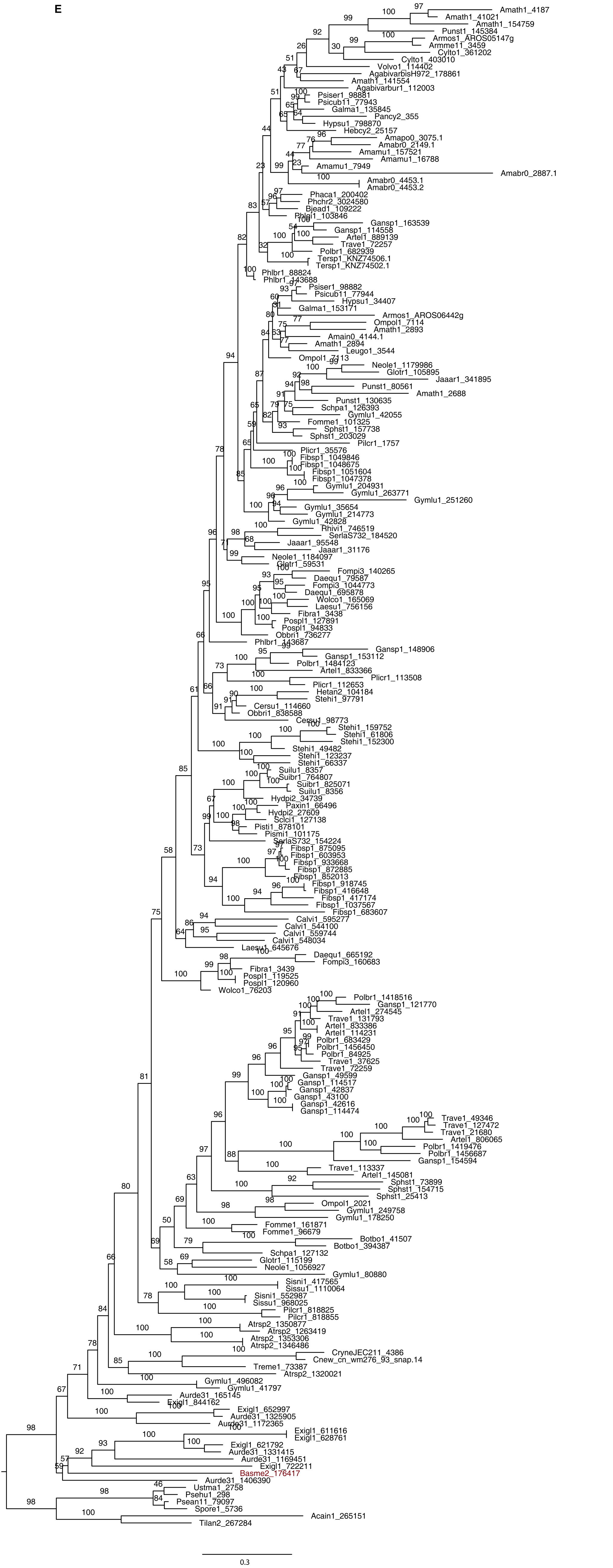

F

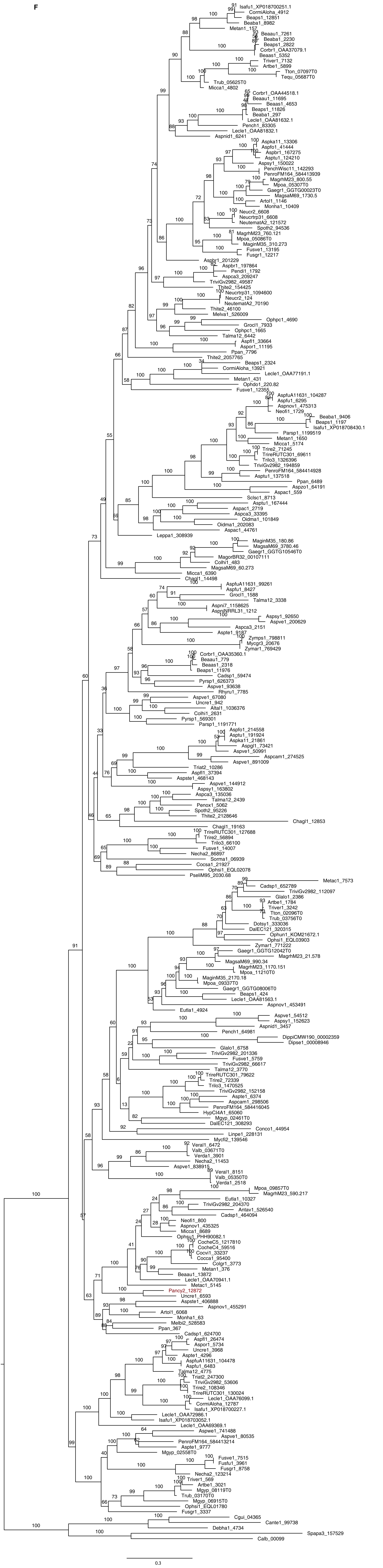

Isafu1_XP018700251.1
CormiAloha_4912
Beaps1_12851
Beaba1_8982
Metan1_157
Beaau1_7261
Beaba1_2230
Beaps1_2822
Corbr1_OAA37079.1
Beaas1_5352
Triver1_7132
Artbe1_5899
Tton_07097T0
Tequ_05687T0
Trub_05625T0
Micca1_4802
Corbr1_OAA44518.1
Beaau1_11695
Beaas1_4653
Beaps1_11826
Beaba1_297
Lecle1_OAA81632.1
Pench1_83305
Lecle1_OAA81832.1
Aspnid1_6241
Aspka11_13306
Aspfo1_41444
Aspbr1_167275
Asptu1_124210
Aspsy1_150022
PenchWisc11_142293
PenroFM164_584413939
MagrhM23_800.55
Mpoa_05307T0
Gaegr1_GGTG00023T0
MagsaM69_1730.5
Artol1_1146
Monha1_10409
Neucr2_6608
Neucrtrp31_6608
NeutematA2_121572
Spoth2_94536
MagrhM23_760.121
Mpoa_05086T0
MaginM35_310.273
Fusve1_13195
Fusgr1_12217
Aspbr1_201229
Aspbr1_197864
Pendi1_1792
Aspca3_209247
TriviGv2982_49587
Thite2_154425
Neucrtrp31_1094600
Neucr2_124
NeutematA2_70190
Thite2_46100
Melva1_526009
Ophpc1_4690
Grocl1_7933
Ophpc1_1665
Talma12_6442
Aspfl1_33664
Aspor1_11195
Ppan_7796
Thite2_2057765
Beaps1_2324
CormiAloha_13921
Lecle1_OAA77191.1
Metan1_431
Ophdo1_220.82
Fusve1_12355
AspfuA11631_104287
Aspfu1_6295
Aspnov1_475313
Neofi1_1729
Beaba1_9406
Beaps1_1197
Isafu1_XP018708430.1
Parsp1_1199519
Metan1_1650
Micca1_5174
Trire2_71245
TrireRUTC301_69611
Trilo3_1326396
TriviGv2982_194859
PenroFM164_584414928
Asptu1_137518
Ppan_6489
Aspzo1_64191
Aspac1_559
Sclsc1_8713
Asptu1_167444
Aspac1_2719
Aspca3_33395
Oidma1_101849
Oidma1_202083
Aspac1_44761
Leppa1_308939
MaginM35_180.86
MagsaM69_3780.46
Gaegr1_GGTG10546T0
MagorBR32_00107111
Colhi1_483
MagsaM69_60.273
Micca1_6390
Chagl1_14498
AspfuA11631_99261
Aspfu1_8427
Grocl1_1588
Talma12_3338
Aspni7_1158625
AspniNRRL31_1212
Aspsy1_92650
Aspve1_200629
Aspca3_2151
Aspte1_9187
Zymps1_798811
Mycgr3_20676
Zymar1_769429
Corbr1_OAA35360.1
Beaau1_779
Beaas1_2318
Beaps1_11976
Cadsp1_59474
Pyrsp1_626373
Aspve1_93638
Rhyru1_7785
Aspve1_67080
Uncre1_942
Altal1_1036376
Colhi1_2631
Pyrsp1_569301
Parsp1_1191771
Aspfo1_214558
Asptu1_191924
Aspka11_21861
Aspgl1_73421
Aspve1_50991
Aspcam1_274525
Aspve1_891009
Triat2_10286
Aspfl1_37394
Aspste1_468143
Aspve1_144912
Aspsy1_163802
Aspca3_135036
Talma12_2439
Penox1_5062
Spoth2_95226
Thite2_2128646
Chagl1_12853
Chagl1_19163
TrireRUTC301_127688
Trire2_56894
Trilo3_66100
Fusve1_14007
Necha2_86897
Sorma1_06939
Cocsa1_21927
Ophsi1_EQL02078
PseliM95_2030.68
Metac1_7573
Cadsp1_652789
TriviGv2982_112097
Glalo1_2386
Artbe1_1784
Triver1_3242
Tton_02096T0
Trub_03756T0
Dotsy1_333036
DalEC121_320315
Ophun1_KOM21672.1
Ophsi1_EQL03903
Zymar1_771222
Gaegr1_GGTG12042T0
MagrhM23_21.578
MagsaM69_990.34
MagrhM23_1170.151
Mpoa_11210T0
MaginM35_2170.18
Mpoa_09337T0
Gaegr1_GGTG08006T0
Beaps1_424
Lecle1_OAA81563.1
Aspnov1_453491
Eutla1_4924
Aspve1_54512
Aspsy1_152623
Aspnid1_3457
Pench1_64981
DippiCMW190_00002359
Dipse1_00008946
Glalo1_6758
TriviGv2982_201336
Fusve1_5759
TriviGv2982_66617
Talma12_3770
TrireRUTC301_79622
Trire2_72339
Trilo3_1470525
TriviGv2982_152158
Aspte1_6374
Aspcam1_298506
PenroFM164_584416045
HypCl4A1_65060
Mgyp_02461T0
DalEC121_308293
Conco1_44954
Linpe1_228131
Mycfi2_139546
Veral1_6472
Valb_03671T0
Verda1_3901
Necha2_11453
Aspve1_838915
Veral1_8151
Valb_05350T0
Verda1_2518
Mpoa_09857T0
MagrhM23_590.217
Eutla1_10327
TriviGv2982_204370
Antav1_526540
Cadsp1_464094
Neofi1_800
Aspnov1_435325
Micca1_8689
Ophsu1_PHH90082.1
CocheC5_1217810
CocheC4_59516
Cocvi1_33237
Cocca1_95400
Colgr1_3773
Metan1_376
Beaau1_13872
Lecle1_OAA70941.1
Metac1_5145
Pancy2_12872
Uncre1_6593
Aspste1_406888
Aspnov1_455291
Artol1_6068
Monha1_63
Melbi2_528583
Ppan_367
Cadsp1_624700
Aspfl1_26474
Aspor1_5734
Uncre1_3968
Aspte1_4296
AspfuA11631_104478
Aspfu1_6483
Talma12_4775
Triat2_247300
TriviGv2982_53606
Trire2_108346
TrireRUTC301_130024
Lecle1_OAA76099.1
CormiAloha_12787
Isafu1_XP018700227.1
Lecle1_OAA72986.1
Isafu1_XP018703052.1
Lecle1_OAA69369.1
Aspwe1_741488
Aspve1_80535
PenroFM164_584413214
Aspte1_9777
Mgyp_02558T0
Fusve1_7515
Fusfu1_3961
Fusgr1_8758
Necha2_123214
Triver1_569
Artbe1_3021
Mgyp_08119T0
Trub_03170T0
Mgyp_06915T0
Ophsi1_EQL01780
Fusgr1_3337
Cgui_04365
Cante1_99738
Debha1_4734
Spapa3_157529
Calb_00099
0.3

**Supplemental Figure 2. Phylogeny of LysM domains**
Maximum Likelihood phylogeny (IQ-tree) of LysM domains extracted from all fungal chitinases. Support values indicate percentage of rapid bootstraps

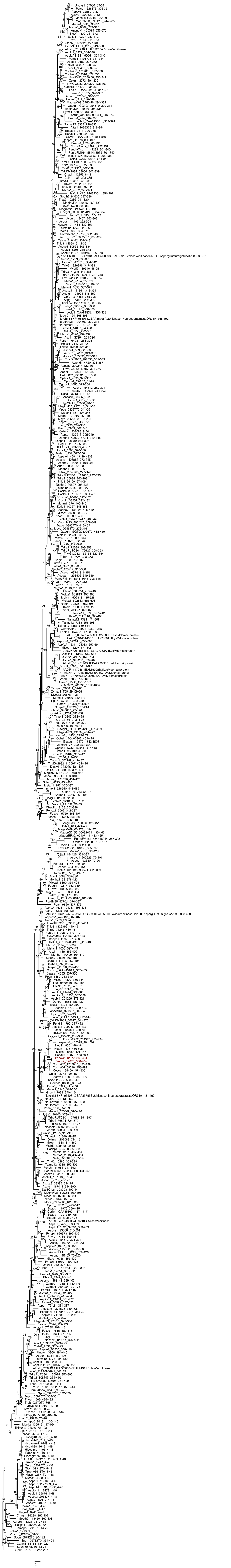
